# Life-long behavioral screen reveals an architecture of vertebrate aging

**DOI:** 10.1101/2025.11.21.688112

**Authors:** Claire N. Bedbrook, Ravi D. Nath, Libby Zhang, Scott W. Linderman, Anne Brunet, Karl Deisseroth

## Abstract

Mapping behavior of individual vertebrate animals across lifespan is challenging, but if achieved, could provide an unprecedented view into the life-long process of aging. We created the first platform for high-resolution continuous behavioral tracking of a vertebrate animal across natural lifespan from adolescence to death—here, of the African killifish. This behavioral screen revealed that animals follow distinct individual aging trajectories. The behaviors of long-lived animals differed markedly from those of short-lived animals, even relatively early in life, and were linked to organ-specific transcriptomic shifts. Machine learning models accurately predicted age and even forecasted an individual’s future lifespan, given only behavior at a young age. Finally, we found that animals progressed through adulthood in a sequence of stable and stereotyped behavioral stages with abrupt transitions suggesting a novel structure for the architecture of vertebrate aging.

## Main text

Deep insight into the processes of life has come from the dynamic observation of intact biological systems over long timescales. For example, continuous inspection of developing embryos has provided critical insight into processes governing the transition from a single cell to a fully formed organism (*1–7*). But these dynamical maps have typically ended with completion of initial organismal development. Fundamentally distinct classes of insight might be derived from continuing to observe individual organisms across the entirety of adulthood all the way to aging-related death. Continuous tracking during aging could reveal new information regarding unrecognized post-maturity life stages, as well as natural health decline during aging, especially as age is the primary risk factor for most chronic diseases (*8–10*). However, given the long timescale of vertebrate aging, such continuous observation across an individual vertebrate animal’s life has not been practical.

Behavior is a rich and diverse readout of animal state (*11–15*)—integrating diverse features of sensation, cognition, and action—which has been shown to reflect the aging process in worms (*16–20*), flies (*21, 22*), mice (*23, 24*), and humans (*25–28*). Recent advances in computer vision and animal-tracking technology (*14, 29–33*) have enabled the continuous recording and quantification of naturalistic, self-motivated behaviors. Yet the required timescale for widely-applied vertebrate model systems (e.g., 2–3 years for median lifespan in mice and ∼3 years median lifespan in zebrafish (*9*)) is prohibitively long for whole-lifespan, continuous recording of behavior. Here, we leverage the African turquoise killifish, a vertebrate model for aging with a naturally compressed lifespan (median 4–8 months) (*34–49*), to continuously record behavior from puberty to death and explore the architecture of adult development and aging. With this life-long behavioral screen, we sought to systematically determine how naturalistic behaviors change with age and whether behavior was sufficient to predict aging differences and even remaining life. Our unbiased approach also allowed us to explore the impact of interventions on behavior and to uncover discrete behavioral stages throughout adult life.

### System for continuous behavioral tracking across life

To continuously measure behavior of individuals in a highly-scalable manner, we designed a system to house and track many individual animals in a physiologically-suitable freely-moving environment over the entire adult lifespan (Fig. 1; fig S1). We began with male African turquoise killifish individually housed with circulating water flow (Fig. 1A). The animals received seven feedings each day at fixed times using custom programmable automated feeders (*50*). Cameras were mounted overhead, and ran continuously at 20 frames per second (Fig. 1B; Movie S1).

**Fig. 1.**
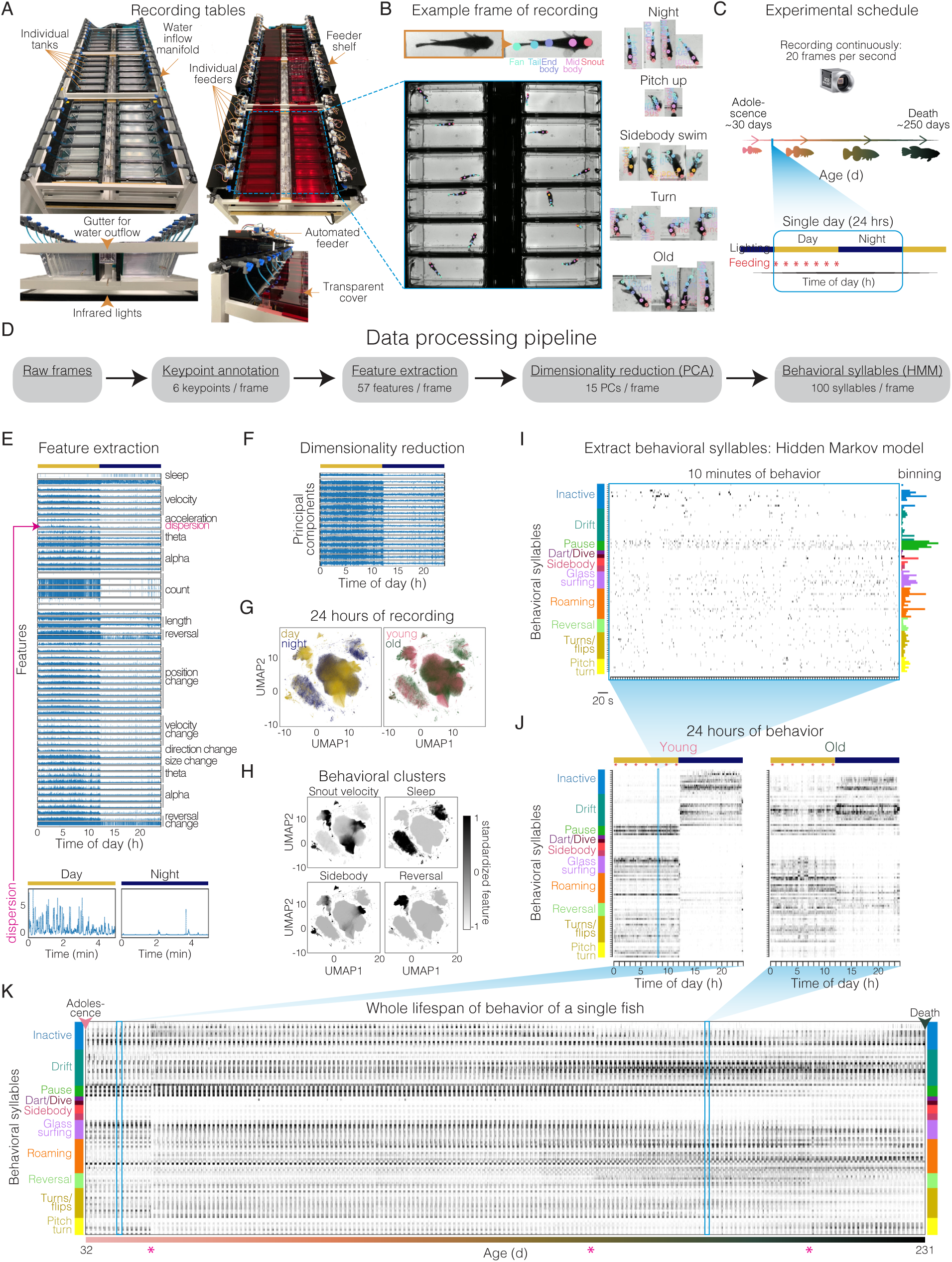
System enables continuous behavioral tracking from adolescence to death. (**A**) Long-term recording setup: 2.8-L tanks fixed in a custom table rack with continuous water inflow and outflow. Automated feeders were fixed above each individual tank on feeder shelf. Infrared-transparent red acrylic lid was placed above all tanks, with infrared backlight below tanks. (**B**) Example frame from Basal industrial camera (capturing 20 frames per second) imaging a single recording table. Six key points (snout, midbody, endbody, tail, fan, and sidebody) were predicted from trained deep convolutional neural network overlayed on example frame. Cropped individual frames of recording show animals at different times of day, age, and positions. (**C**) Experimental schedule for recordings from adolescence to death with 12-hr/12-hr light/dark cycle. Yellow indicates day, and blue indicates night. (**D**) Data-processing pipeline from raw video frames to behavioral syllable usage. (**E**) Pose features extracted from key point data for a single animal in one day (24-hr cycle) of its life. Inset below shows zoom-in of 5-min interval during the day and night highlighting dispersion. (**F**) PCA of pose features for same animal and day from data shown in **E**. (**G**) UMAP embedding of pose features PCs of data from a single day for nine different fish down sampled to show one every five frames, with each point being a single frame of recording and color maps indicating day vs. night frames or young (45 d) vs. old (270 d) frames. (**H**) UMAP embedding of pose features PCs with heatmap for four select pose features. (**I**) Sampling of 100 HMM-derived behavioral syllables for a single fish during 10-min interval during one day of its life. Behavioral syllables are ordered according to similarity (Kullback–Leibler divergence [D_KL_]) and hierarchical clustering of distinct types of behavior shown in color along left (see fig. S4B for clustering). Histogram shows time spent in each behavioral syllable during the 10-min bin on right. (**J**) Behavioral syllable usage across 24 hr for a single animal when young (40 d) compared to 24 hr when same animal is old (180 d). (**K**) Whole-lifespan behavioral syllable usage for single animal from 32 days old until death at 231 days with pink asterisk highlighting transitions in syllable use (see Fig. 6 for analysis of these transitions across the population).

Each animal was placed into its recording tank at puberty (sexual maturity, 3–4 weeks of age) and all behaviors were tracked continuously until death (Fig. 1C).

To track animal movement, we trained a deep convolutional neural network that predicts six key points along the body in all frames of the whole-lifespan recording (Fig. 1B; Movie S2; see Methods). From tracked key points for each animal, we calculated ‘pose features’ (a total of 57; Fig. 1D; Table S1) (see Methods) which describe parameters such as velocity, body curvature, and swim direction (Fig. 1E). Followed by principal component (PC) analysis (PCA) to represent pose feature dynamics with fewer dimensions (57 pose features to 15 PCs with >80% variance explained; Fig. 1F, fig. S2A). Pose features over a 24-hour interval revealed distinct behavioral clusters (Fig. 1G-H; fig. S2B; fig. S3). Different behavioral clusters were used in day versus night (linear separability mean cross validation accuracy=0.71) (Fig. 1G; fig. S2C). In contrast, young (45 d) and aged (270 d) animals showed a large degree of overlap in behavioral cluster usage (linear separability mean cross validation accuracy=0.56) (Fig. 1G; fig. S2D; fig. S3), indicating that aging did not completely restructure the behavioral repertoire.

To investigate how the behavioral repertoire did change across lifespan in individual animals, we extracted behavioral motifs. We took an unsupervised approach using a Gaussian hidden Markov model (HMM) to model stereotyped sub-second behavioral motifs from the pose feature dynamics, referred to as “behavioral syllables” (Fig. 1I-K). HMM-derived behavioral syllables (*11, 13, 51*) have been shown to reflect animal internal states (*52–54*). Due to the unprecedented amount of data, we developed a distributed, stochastic expectation-maximization procedure to fit the HMM to >3×10^8^ frames of data (fig. S4A) (see Methods), resulting in 100 behavioral syllables including active behavioral syllables (e.g., flips, bursts, and reversals) as well as inactive behavioral syllables (e.g., slow drifting and resting). Over 24 hours, animals sampled distinct behavioral syllables during the day and night (Fig. 1J; fig. S5A). We observed a clear shift in behavioral syllable usage for individual animals at young (40 d) versus aged (180 d) timepoints (Fig. 1J). Continuous sampling of behavior across the whole adult lifespan of individual animals enabled the first investigation of vertebrate multidimensional behavioral dynamics across timescales from frames (milliseconds) to whole lifespans (∼250 days from puberty to death) in a systematic and quantitative manner (Fig. 1K; fig. S6).

### Global behavioral changes during aging from whole-lifespan tracking

To identify behavioral changes during aging that were consistent across many individual animals, we developed an approach to address the challenge of massive dimensionality of the dataset. For each animal on each day, there are 1.728×10^6^ frames of recording with one of 100 possible behavioral syllables per frame, and for lifespan recordings 81 animals were recorded for a mean of 220 days (range: 23 d to 371 d). To represent daily behaviors by fewer dimensions while preserving the strong circadian structure within the dataset, we used tensor component analysis (TCA) (*55*), which is an unsupervised method capable of extracting low-dimensional factors from multi-dimensional data. We trained nonnegative TCA models on the whole-lifespan timeseries restructured into daily behavioral syllable usage (binned at 10 min resolution) concatenating animals across the time dimension such that a single TCA model was trained for all fish (Fig. 2A). Increasing the number of tensor components (TCs) improves reconstruction error but improvement becomes marginal by ∼45 TCs for the held-out test set (∼40% unexplained variance; Fig. 2B), and reconstructions built using a 45-TC model strongly resemble raw data (Fig. 2C); we therefore used a model with 45 TCs for next-stage analysis.

**Fig. 2.**
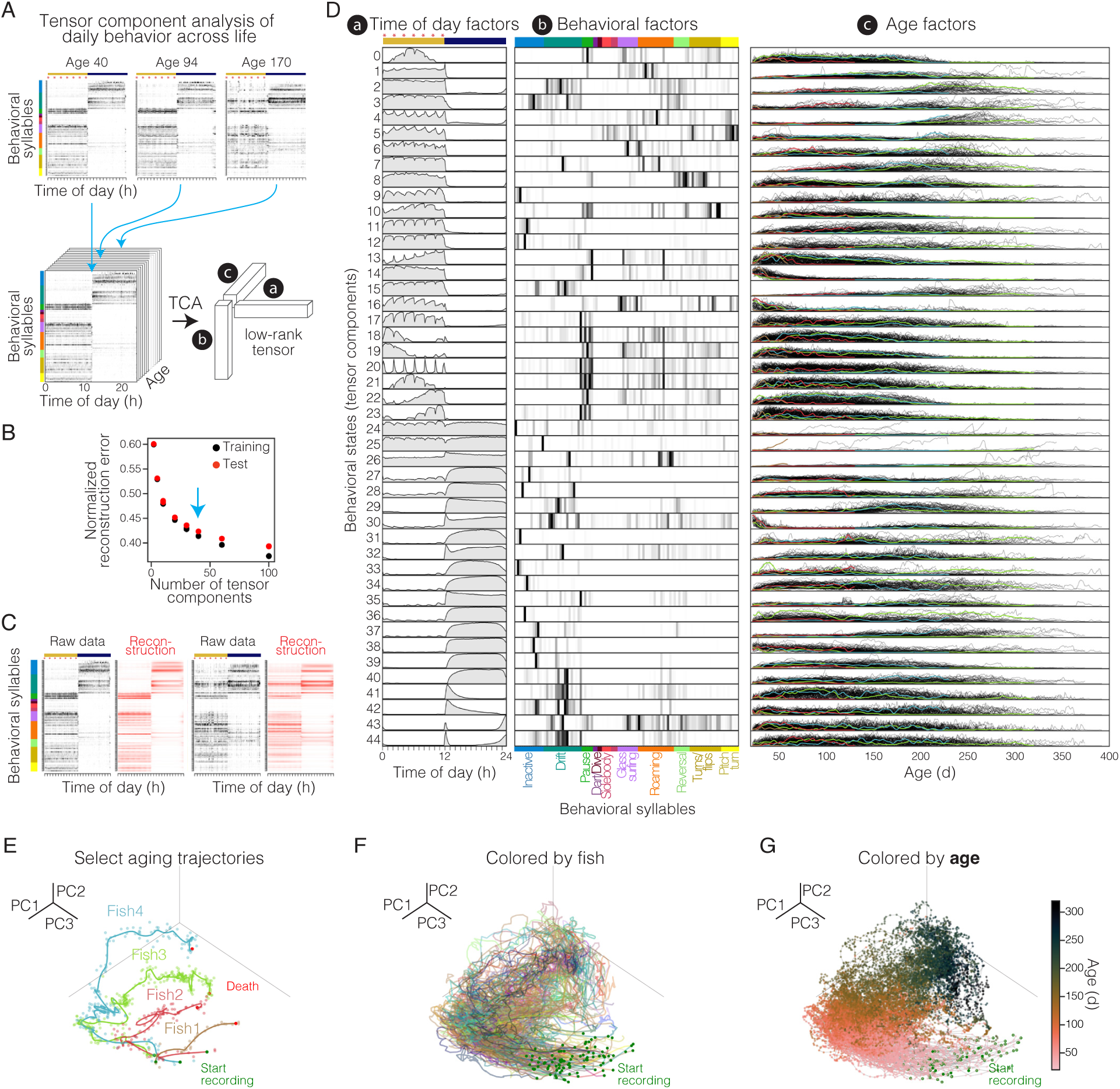
Global behavior changes with age reveal distinct aging trajectories. (**A**) Schematic of TCA setup to investigate whole-lifespan behavioral syllable use. Throughout figure, red asterisks indicate feeding times; light/dark cycle is indicated by yellow/blue bar; behavioral syllables are ordered and clustered as in Fig. 1I. (**B**) Normalized reconstruction error for both training and held-out test set data for models trained with different numbers of TCs. (**C**) Comparison of raw vs. TCA reconstructed syllable data for two randomly selected days. (**D**) Trained 45-TC model showing time-of-day factor, behavioral-syllable factor, and age factors for each TC (i.e., behavioral state). Age factors show weighting of each factor over age, with each thin line representing the trajectory of a single animal. Select animals shown in color correspond to animals shown in **E**. (**E**) Top three PCs of 45 TC data to visualize aging trajectories of select individual animals from start of recording (green circle) to death (red circle). Each animal is shown in a different color. Points show individual days. Line shows smoothed trajectory. Aging trajectories of >100 animals, with each animal in a different color (**F**) and with each point colored by animal age (**G**).

TCA approximates the data as a sum of outer products of three vectors—in our case: a) time-of-day factors, b) behavioral-syllable factors and c) age factors (Fig. 2A, D). The time-of-day factors recapitulate the expected circadian dynamics, including distinct TCs dominant during the day (e.g., TC1) or night (e.g., TC29), as well as TCs that peak at each of the seven feedings (e.g., TC20) (Fig. 2D, subpanel a). This analysis groups behavioral syllables commonly used together with specific temporal dynamics throughout the circadian cycle to build more complex behavioral states, providing a structure for comparison across ages. Indeed, the amplitude or weighting of each time-of-day and behavioral-syllable factor differs each day and for each animal, and is represented as the third factor, the ‘age factor’ (Fig. 2D, subpanel c). While many TCs show changes with age (e.g., TC14), others show more uniform use across age (e.g., TC44), and yet other TCs (e.g., TC24) show large increases in use with age but in only a small subset of animals (Fig. 2D, subpanel c). This TCA analysis provides a robust way to reduce data dimensionality, enabling both qualitative and quantitative comparisons between behavioral states across lifespan.

### Distinct aging trajectories based on behavior

Do individuals exhibit distinct behavioral aging trajectories? To investigate this, we visualized age factors across life for each animal. We performed PCA on all 45 TCs identified above and plotted the top three PCs (Fig. 2E-G) (PC1–3: 40% explained variance; fig. S7). Focusing on the aging trajectories of select fish illustrates that individual animals exhibited distinct aging trajectories in behavioral space (Fig. 2E), suggesting that individuals are not all aging in the same way. The behavioral trajectories are not linear: animals spent more days in some parts of behavioral space than others (Fig. 2E; fig. S8-S9). Interestingly, across large cohorts of animals, young animals generally started in a similar place (low variance across 45 TCs; fig. S10A), but we observed that aging animals fanned out into distinct behavioral trajectories (Fig. 2F) (increasing variance with age across all 45 TCs; fig. S10A), and the position along these behavioral trajectories was strongly correlated with animal age (Fig. 2G). We obtained similar results using UMAP embedding for visualization instead of restricting observations to the top three PC dimensions (fig. S10B-C).

Given that behavior is a non-invasive and non-destructive readout, we recorded across the complete adult lifespan of each individual. There was variability in lifespan, from short (46 d) to long (394 d) (Fig. 3A), as is typical of all species even in genetically identical populations (*56*). Marking trajectories by the *lifespan* of each fish revealed that animals with a long lifespan (Fig. 3B; blue) exhibited a distinct trajectory from that of animals with a short lifespan (Fig. 3B; yellow). Furthermore, comparing the mean trajectories of animals with long vs. short lifespan revealed that long-lived (≥200 days) and short-lived (≥120 days and <200 days) trajectories started together in youth but diverged quite early in life with a branch point at ∼70 days old (Fig. 3B). Interestingly, this branching took place when animals were still relatively young and a long time before death, even for the short-lived population (50–130 days prior to death), suggesting physiological changes relatively early in life that dictate an animal’s aging trajectory and future lifespan, and similarly for healthspan (fig. S11).

**Fig. 3.**
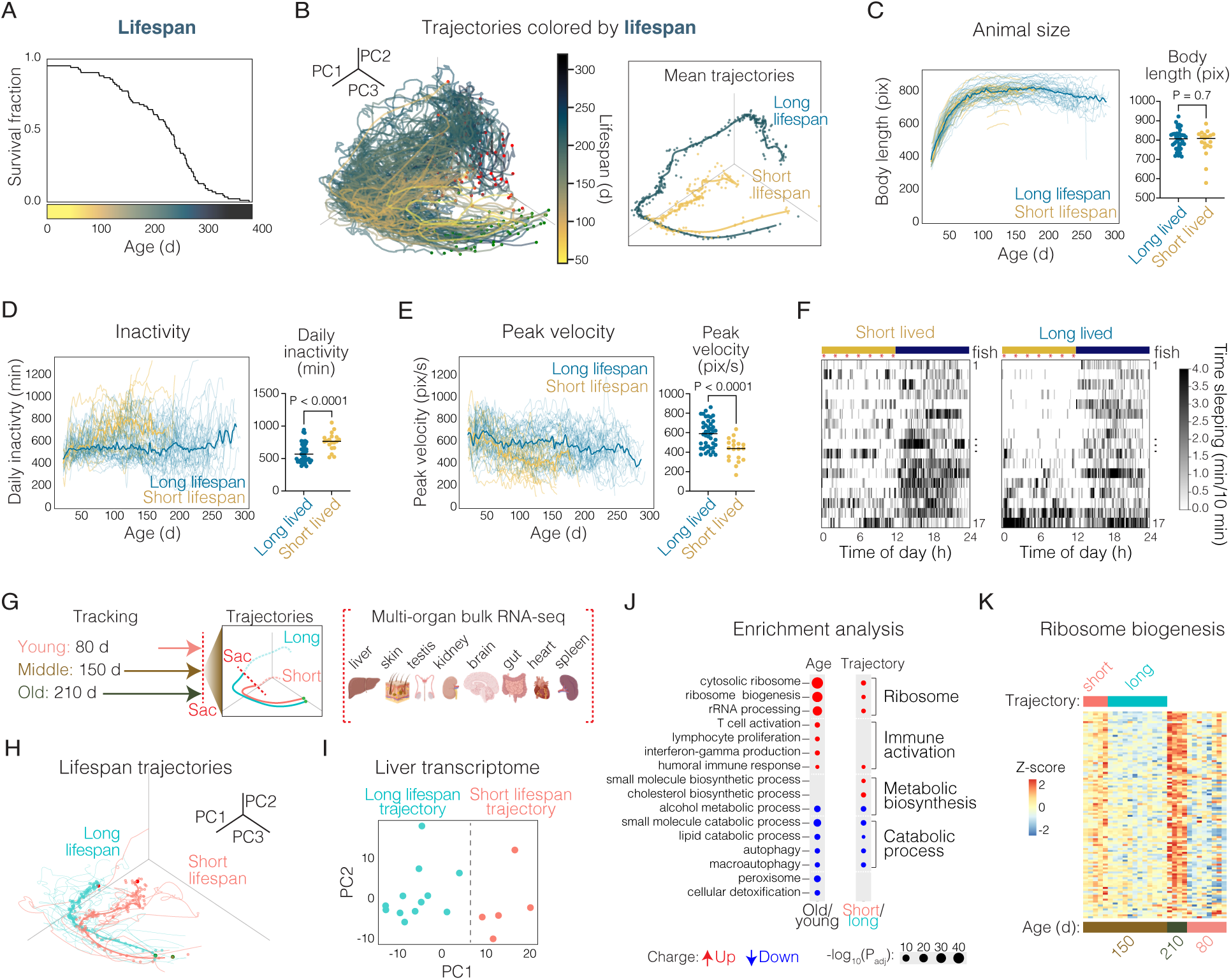
Behavioral aging trajectories diverge early in life based on future lifespan and are linked to large transcriptomic changes in the liver. (**A**) Kaplan–Meier lifespan curve of tracked animals (n=81). (**B**) Aging trajectories of 81 animals colored by future lifespan. Inset shows mean trajectory of distinct lifespan groups: long lifespan (≥ 200 days) and short lifespan (<200 days and ≥120 days to exclude the impact of individuals that died prematurely). Animal size (length) (**C**), inactivity (**D**), and peak velocity (**E**) across life (left) and at 100 days old (right) grouped by lifespan. Mann Whitney test for significance. At 100 days old, there is a significant positive correlation between peak velocity and future lifespan (Spearman’s correlation: P=0.005, π=0.35). (**F**) Heatmap of time spent sleeping across 24 hr at 100 days old with each row representing a different animal and animals grouped by lifespan. Red asterisks indicate feeding times. Light/dark cycle is indicated by yellow/blue bar. Seventeen animals were randomly sampled from the long-lived group to match the size of the short-lived group. (**G**) Scheme for organ RNA-sequencing experiment of animals tracked to different age endpoints, young (80 d; n=8), middle-aged (150 d; n=17), and old (≥210 d, n=4). At each endpoint, organs were harvested for bulk RNA-sequencing. (**H**) Behavioral aging trajectories of 17 animals colored by inferred future-lifespan trajectory. (**I**) PCA of liver transcriptome for each animal in the middle-aged group (150 d; n=17) colored by future-lifespan trajectory based on behavior. (**J**) Gene ontology (GO) enrichment analysis of the liver transcriptome as a function of age (old vs. young) and lifespan trajectories based on behavior (short-vs. long-lifespan trajectory) highlighting significant GO terms. (**K**) Heatmap of genes involved in ribosome biogenesis GO term. Each row is a gene and each column is an animal. Color bar below heatmap indicates the age-group of each individual animal and the color bar above shows the lifespan trajectory of the middle-aged (150 d) group.

What is different between long-vs. short-lived animals? There were not large differences in animal size (length) (Fig. 3C), growth rate, or feeding behavior (tensor component #20 in Fig. 2D; fig. S12B) in the long-vs. short-lifespan groups across life; moreover, both long-lived and short-lived animals spent a similar proportion of their life prior to death with reduced food consumption resulting in residual uneaten food (fig. S12A). Thus, multiple quantitative measures revealed no evidence for dramatic differences in food consumption between the two groups.

Long-lived and short-lived animals also showed similar peak and median velocity at young ages (<70 days). However, by 100 days, long-lived animals spent significantly more time active than short-lived animals (P<0.0001, Mann Whitney test) and exhibited significantly higher peak velocity (“sprinting”) and median velocity (peak velocity, P<0.0001; median velocity, P<0.0001 Mann Whitney test) (Fig. 3D-E; fig S12D). Consistent with this finding, long-lived animals maintained more elevated vigorous darting behavior levels compared with short-lived animals by 100 days (P<0.0003 Mann Whitney test; fig S12C). In addition to these differences in activity and velocity, we also observed dramatic differences in the circadian timing of active and inactive behaviors, comparing the long-and short-lived groups at 100 days (Fig. 3F; fig S12F).

Specifically, short-lived animals showed increased sleep behavior during the light/day period while sleep behavior was more restricted to the dark/night period in the long-lived group at age-matched timepoints. Thus, prior to middle age, animals destined for a long lifespan (while the same size) were more active, had higher peak velocity, and showed tighter circadian timing of sleep and active behaviors.

### Molecular differences between long-lived and short-lived aging trajectories

The continuous behavioral tracking system was designed to non-invasively evaluate the aging trajectory of animals while still relatively young, allowing comparison of animals destined for a long lifespan to those destined for a short lifespan even while early in life. We tracked a cohort of animals from adolescence to middle age (150 days old; n=17 animals) at which point we could reliably distinguish long-lifespan vs. short-lifespan trajectories based on behavior for each individual (Fig. 3G-H). We then sacrificed animals and performed bulk RNA sequencing of eight different organs (brain, gut, heart, kidney, liver, skin, spleen, and testis) of each individual to investigate molecular signatures of distinct aging trajectories. For context, in addition to evaluating animals on a long-vs. short-lifespan trajectory, we also compared molecular signatures of young vs. old animals. For this measurement, we tracked a cohort of animals up to a relatively young age (80 days; n=8) and a cohort of animals up to advanced age (≥210 days; n=4) (Fig. 3G); both groups were sacrificed followed by bulk RNA sequencing of the eight different organs.

Across multiple organs, PCA could separate the transcriptomes of individuals of the same chronological age but of different lifespan trajectories based on behavior (fig. S13A-B).

Interestingly, the liver had the clearest separation of individual transcriptomes associated with long-lifespan or short-lifespan trajectory based on behavior (Fig. 3I), as quantified by Silhouette score (fig. S13B). Unbiased k-means clustering of the liver transcriptomes resulted in two clusters that matched the short-lifespan and long-lifespan trajectory groups (fig. S13C), corroborating liver transcriptome separation. This result was intriguing in light of the liver’s key role in metabolism and sensitivity to interventions that impact longevity (*57–59*), suggesting possible metabolic underpinnings for the differences in behavioral activity and vigor of long-lived vs. short-lived animals.

Gene set enrichment analysis comparing the liver transcriptomes of individuals on long-vs. short-lifespan trajectories revealed major differences in gene ontology (GO)-terms relating to ribosome biogenesis, rRNA processing, translation and DNA replication, all upregulated in animals on a short-lifespan trajectory (Fig. 3J; fig. S14B; Table S2). Interestingly, these same terms were all found to be also significantly upregulated with age (Fig. 3J; fig. S14B). By contrast, several age-related changes were not different in animals on short-vs. long-lifespan trajectories. For example, we also observed a robust increase with age in GO terms relating to immune activation (T-cell activation, lymphocyte proliferation, interferon-gamma production) (Fig. 3J), but this result was not seen when comparing animals on short-vs. long-lifespan trajectories. Thus, immune or inflammation changes may not causally underlie the differences in lifespan we observe but rather represent a phenotype associated with natural aging. By contrast, ribosome biogenesis upregulation (as well as downregulation of catabolic processes such as autophagy; Fig. 3J), seen both in old animals and in animals destined for short lifespan based on behavior (Fig. 3K), could represent candidate molecular drivers of longevity.

### Behavior suffices to predict age

We asked if behavioral measures could suffice to predict the age of an individual—which would be a key step for both quantifying aging and predicting the effect of interventions (*60–62*), especially since behavior is a noninvasive readout well-suited for work across the lifespan. To address this question, we built a behavioral clock—i.e., a statistical model that takes as its input the 45-dimensional age factors (Fig. 2D, subpanel c) that summarize a fish’s behavior for a given day (i.e., daily ‘behaviorome’) in an animal’s life, and predicts the age of the animal on that day (Fig. 4A). We trained a random forest regressor and evaluated model performance using leave-one-fish-out cross validation. The age predicted by this model was found to correlate strongly with true age (*R* = 0.94; Fig. 4B), with a median absolute error (MAE) of only 12 days across ages. At very old ages (>250 days old) the MAE increased to >20 days, likely because of the large drop-off in the amount of training data at these extreme ages (Fig. 4C). We observe consistent results when evaluating model performance on a held-out test cohort of animals (*R* = 0.92, MAE 12 days across ages; fig. S15). These results indicate that behavior of an individual from noninvasive recording is sufficient to closely predict animal age across lifespan.

**Fig. 4.**
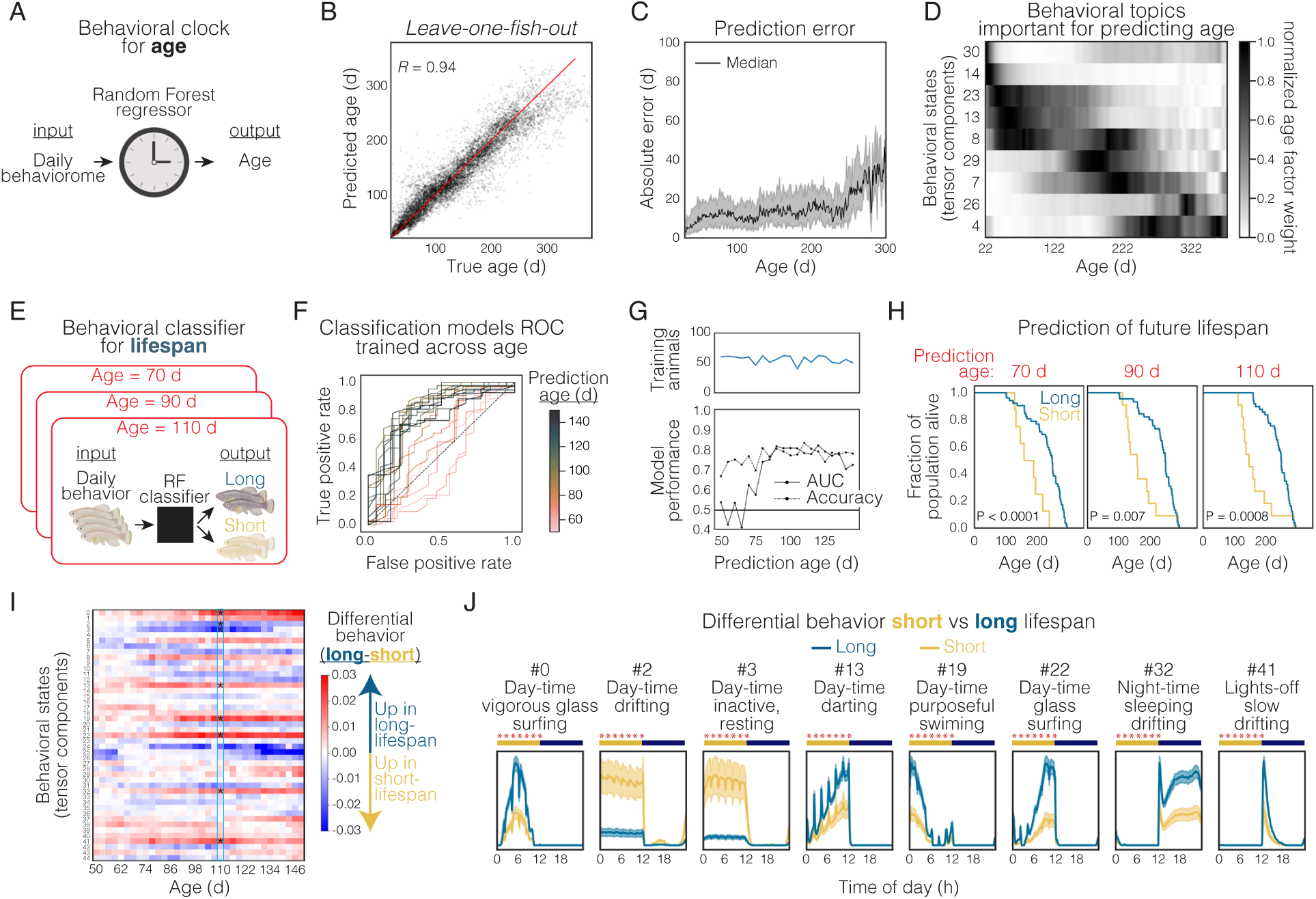
Behavior is highly predictive of age and future lifespan. (**A**) Random forest regression models trained on the 45-dimensional age factors that summarize a fish’s behavior for a given day (i.e., daily ‘behaviorome’) to predict age (i.e., behavioral clock for age). (**B**) Leave-one-fish-out model predictions vs. true age. (**C**) Leave-one-fish-out model prediction absolute error calculated across age. (**D**) Age weighting of daily behavioral states (tensor components) most important for model prediction (based on mean decrease in impurity). (**E**) Random forest classifiers trained on behavioral TC data at distinct age windows to classify animals as either short-or long-lived (i.e., behavioral classifier for lifespan). (**F**) ROC curve for random forest classification models trained on behavioral data from animals at different ages. Coloring of curves is based on age of the animals used for predictions. (**G**) (top) Number of animals included in model training across different age groups. (bottom) Classification model performance in both area under the ROC curve (AUC) and accuracy across different age groups. (**H**) Kaplan–Meier lifespan curve of animals predicted to be short-lived (yellow) and long-lived (blue) at various prediction ages (Log-rank test). (**I**) Differential behavior of long-lived minus short-lived animals, highlighting specific behavioral states that are significantly different between short-and long-lived groups at the age of 110 d (Mann–Whitney U test with Bonferroni correction). (**J**) Highlighting behavioral states (tensor components) that are significantly different between short-and long-lived groups at 110 d, plotting mean age-factor weights by time-of-day factor for the short-lived (yellow) vs. long-lived (blue) groups (tensor component numbering as in Fig. 2D).

Interestingly, the daily behavioral states most important for model prediction were observed to change as a function of age (Fig. 4D). Specifically, the behavioral states most important to predict age (based on mean decrease in impurity; see Methods) at each stage of life were: i) day-time vigorous swimming and exploring behavior in young animals, ii) night-time slow drifting and sleep behavior in middle-aged animals, iii) constant (noncircadian) slow drifting turns in old animals, and iv) drifting backwards and reversal flips in very old (geriatric) animals. These waves of distinct behavioral state usage at specific age ranges reveal that aging in adult animals may not be well described as a gradual decline but rather as progression through discrete stages in which distinct combinations of behaviors are used.

### Future-lifespan forecasting based on behavior

We next asked if daily behavioral measures could predict an animal’s future lifespan. Specifically, would it be possible to forecast, based solely on behavior at a young age, if animals were destined for short vs. long lifespans? To test this possibility, we built classification models at different fixed age ranges. These models input the 45-dimensional age factors (Fig. 2D, subpanel c) that summarize a fish’s behavior for a given day (i.e., daily ‘behaviorome’) within a specific age window, and then classify the animal as short-lived (<200-d lifespan) or long-lived (Fig. 4E). While classifier performance to predict future lifespan was found to be low at very young ages (<70 d), dramatic improvement was observed after 70 d (young adults) (receiver operating characteristic (ROC) curve and accuracy >0.7; Fig. 4F-G).

Importantly, we used the age-fixed classifiers to classify each animal within a cohort as short-vs. long-lived (leave-one-fish-out predictions) and then plotted the true survival curves for the separate groups. This analysis revealed a large and significant separation in lifespan curves (Fig. 4H). Future lifespan predictions were accurate using behaviors at age 70, 90, and 110 d (all before midlife) (Fig. 4H). Thus, using behavior alone, we could accurately forecast, even at a young age, if individuals were destined to be short-lived or long-lived.

We next asked which specific behaviors, at a young age, might be most associated with individuals destined for a long lifespan vs. a short lifespan. To this end, we quantified differential behavior usage (defined as the difference between age factors in long-lived vs. short-lived animals) over a range of ages from 50 to 150 d (Fig. 4I). Clear behavioral differences were seen between long-lived and short-lived animals even at young ages (Fig. 4I), but the separation became stronger as the animals aged. Focusing on 110 d (before middle-age, and when behavioral classifiers are highly accurate), we observed stark differences in behaviors between the animals that were long-vs. short-lived (Mann–Whitney U test with Bonferroni correction; Fig. 4I-J). Animals destined for a long lifespan exhibited more time in vigorous/active states during the day compared to animals destined for a short lifespan (Fig. 4J). Additionally, individuals destined for a long lifespan, even at a relatively young age, displayed a tighter night sleep pattern, with dramatically less time in sleep/rest states during the day and increased time in inactive drift and sleep states during the night, compared to animals destined for a short lifespan (Fig. 4J). The linkage of long lifespan with well-defined circadian timing of behavior (especially sleep and inactivity-related actions) may be consistent with findings implicating circadian timing of feeding in longevity (*59, 63*).

Taken together, these new datasets and analyses revealed that behavior, even at a relatively young age, can forecast remaining life, with potentially major implications for identifying interventions that impact life trajectories.

### Behavioral changes with a longevity intervention: dietary restriction

Quantitative and non-invasive evaluation of the progression of aging in individuals provides a unique capability to explore how longevity interventions influence the process of aging.

Longevity interventions such as dietary restriction (both calorie-restricted and time-restricted feeding) have been shown to extend lifespan and promote health in animals across the animal kingdom from *C. elegans* to non-human primates (*64, 65*), including in killifish (*50, 66*). We next asked how dietary restriction might influence behavior and progression along aging trajectories? To address these questions, we tracked behavior in a cohort of male individuals (n=39) undergoing a dietary restriction regime that has previously been reported to result in lifespan extension in killifish (*50*), with feedings only 3 times per day and all during the morning (Fig. 5A). Animals undergoing dietary restriction were tracked and behavior analyzed from adolescence until 110 days of age. We compared the behavioral characteristics and aging trajectories of animals undergoing dietary restriction to prior cohorts of animals undergoing *ad libitum* feeding (*ad libitum* reference population with 7-times per day food access spaced evenly throughout the 12-hr light period; Fig. 5A).

**Fig. 5.**
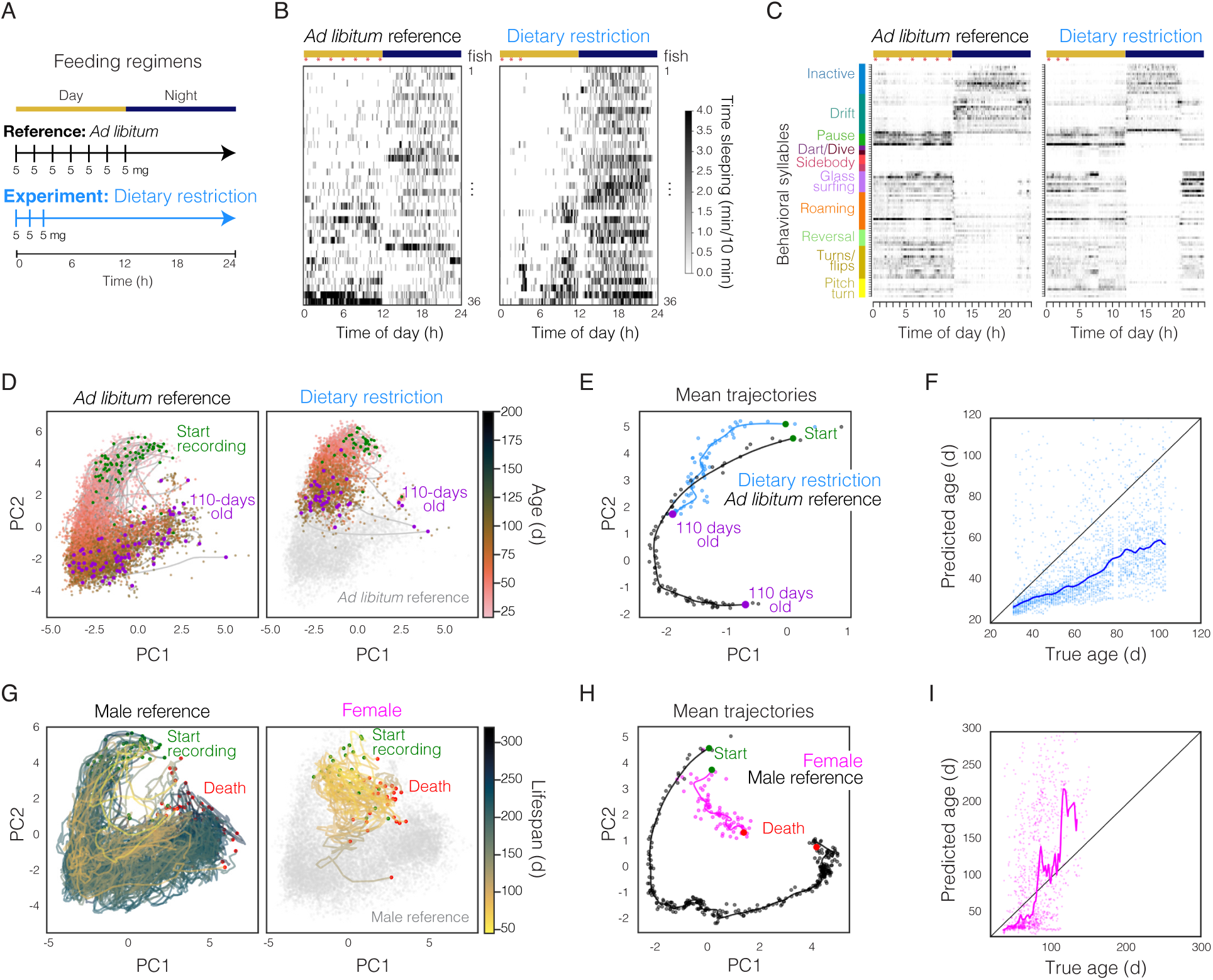
Influence of conserved longevity intervention and sex on behavior and behavioral aging trajectories. (**A**) Scheme comparing the experimental dietary restricted feeding regime with the *ad libitum* reference feeding regime. (**B**) Heatmap of time spent sleeping across 24 hr at 54 days old, with each row representing a different animal. Animals under dietary restriction (right; n=36) are compared to randomly selected *ad libitum* reference animals from previous cohorts (left; n=36). Red asterisks indicate feeding times. Light/dark cycle is indicated by yellow/blue bar. (**C**) Behavioral syllable usage across 24 hr for a single animal under dietary restriction (right) and a single representative animal selected from the *ad libitum* reference cohorts at 33 days old. See fig. S16F for mean behavioral syllable usage across all animals under each feeding regime. Behavioral syllables are ordered and clustered as in Fig. 1I. (**D**) Behavioral aging trajectories of animals under dietary restriction (right; n=39) and animals of the *ad libitum* reference cohorts (left; n=80) with each point colored by animal age. Points represent individual days. Line shows smoothed trajectory for an individual animal. Purple points indicate when animals reach 110 days old. (**E**) Mean aging trajectory of *ad libitum* reference animals (black) vs. dietary restriction animals (blue). (**F**) Behavioral clock model predictions vs. true age for dietary restricted animals. (**G**) Aging trajectories of females (right; n=31) and male reference animals (left; n=118; *ad libitum* feeding) colored by lifespan. Line shows smoothed trajectory for an individual animal. Red points indicate when animals die. (**H**) Mean aging trajectory of male reference animals (black) vs. female animals (magenta). (**I**) Behavioral clock model predictions vs. true age for females.

Animals undergoing this dietary restriction regime demonstrated striking changes to the circadian timing of active and sleep behaviors (Fig. 5B; fig. S16E). Specifically, animals under dietary restriction were found to awaken earlier in the morning (Fig. 5B; fig. S16E) with an increase in active behavioral syllables hours prior to the light change (Fig. 5C; fig. S16F).

Despite their early awakening, animals undergoing dietary restriction in fact exhibited increased total sleep duration (fig. S16C; sleep at 60 days old P=0.01, Mann Whitney test) with sleep timing more restricted to night (dark) hours (Fig. 5B). While animals undergoing dietary restriction revealed no significant differences in median movement velocity (fig. S16A), these animals did exhibit significantly elevated peak velocity relative to the *ad libitum* reference (fig. S16B; peak velocity at 60 days old P<0.0001, Mann Whitney test), aligned with our previous observation that lifespan is positively correlated with peak velocity at a relatively young age.

Interestingly, when we compared the behavioral aging trajectory of animals undergoing dietary restriction until 110 days of age with the *ad libitum* reference population, we observed that the two groups followed a similar aging trajectory based on behavior in PC1 and PC2, but that animals undergoing dietary restriction moved along that trajectory much more slowly (Fig. 5D-E). Indeed, when we quantified individual behavioral age using our behavioral clocks of age, we discovered that animals under dietary restriction were predicted to be behaviorally younger than the true chronological age by 42 d (median error from 100–110 days old) (Fig. 5F), revealing that animals undergoing dietary restriction age in a similar manner but at a reduced rate (fig. S16H; P<0.0001, Mann Whitney test). This finding may be consistent with studies in which calorie restricted animals (mice and non-human primates) die with similar pathologies and diseases as controls but at older ages (*59, 67, 68*). Here we focused tracking analysis to adolescence until 110 days of age, and thus this analysis was terminated before assessment of the impact of dietary restriction on lifespan in this cohort (given the sensitivity of lifespan to slightly different experimental conditions, see Methods). It will be interesting to extend this analysis to test if the trajectory observed continues until death and whether it is linked to lifespan extension.

### Life-long behavioral aging in females

Aging is sexually dimorphic across the animal kingdom (*69*), including in humans (*70, 71*), and it is critical to study the process of aging in both sexes. We therefore were curious to explore if female killifish age in a similar manner to males or instead follow a distinct path through life; with our system, we were well-positioned to address this important question. We tracked behavior in a cohort of female killifish (n=31) from adolescence to natural death. With the feeding paradigm and husbandry conditions used for lifespan experiments, in which animals are individually housed and are not bred throughout life, we observed that females are short-lived compared to males (fig. S17E). We compared the aging trajectories of females to our previously-tracked cohorts of males undergoing matched feeding (*ad libitum*) and tracking conditions (Male reference; Fig. 5G-H). Interestingly, we observed that females progress through a distinct aging trajectory relative to the male reference population (Fig. 5G-H). The female behavioral aging trajectory was found to be similar to the trajectory of short-lived males (Fig. 5G) and differed dramatically from the mean male trajectory (Fig. 5H). When using the behavioral aging clock (trained on male reference data), we observed that at young ages (<50 days old) females were predicted to be behaviorally younger than true chronological age. With aging, however, females exhibited a dramatic acceleration in aging trajectory based on behavior, and females >100 days old were predicted to be 50-200 days older than true chronological age (Fig. 5I). Moreover, most (74%) of the female animals were correctly predicted to be short lived based on the behavioral classifier for lifespan at 70 days of age (fig. S17F), suggesting that some (but not all) of the behaviors that are important for longevity model prediction are consistent between females and males. Thus, our behavioral classifier for lifespan (trained on a male population with only 33% short-lived individuals) was nevertheless able to predict greater than twofold enrichment of short-lived females. These results suggest that this model already substantially generalizes despite the large differences between male and female aging, although this model will be extended in the future with training data from both sexes. Notably, females exhibited significantly lower median locomotor velocity and more time sleeping than males (but did not exhibit different peak velocity) (fig. S17A-C); females were also smaller in size than males (fig. S17D). Overall, we discovered that killifish females progress through a distinct and accelerated aging process, consistent with our observation of shorter female lifespan in individually-housed conditions (fig. S17E). These data raise the possibility that female deaths may be associated with distinct causal pathologies compared with males, although further understanding of why females and male killifish age along distinct trajectories remains unresolved.

### Architecture of aging via distinct stereotyped life stages

Continuous readout of aging from adolescence to death in a vertebrate provides the unique opportunity to examine and model the stereotyped progression through adult life. When each individual behavioral syllable was plotted across life with each day circadian phase-aligned, we were surprised to observe multiple discrete, sharp, and stable behavioral transitions across the lifespan (Fig. 6A-B; fig. S18A).

**Fig. 6.**
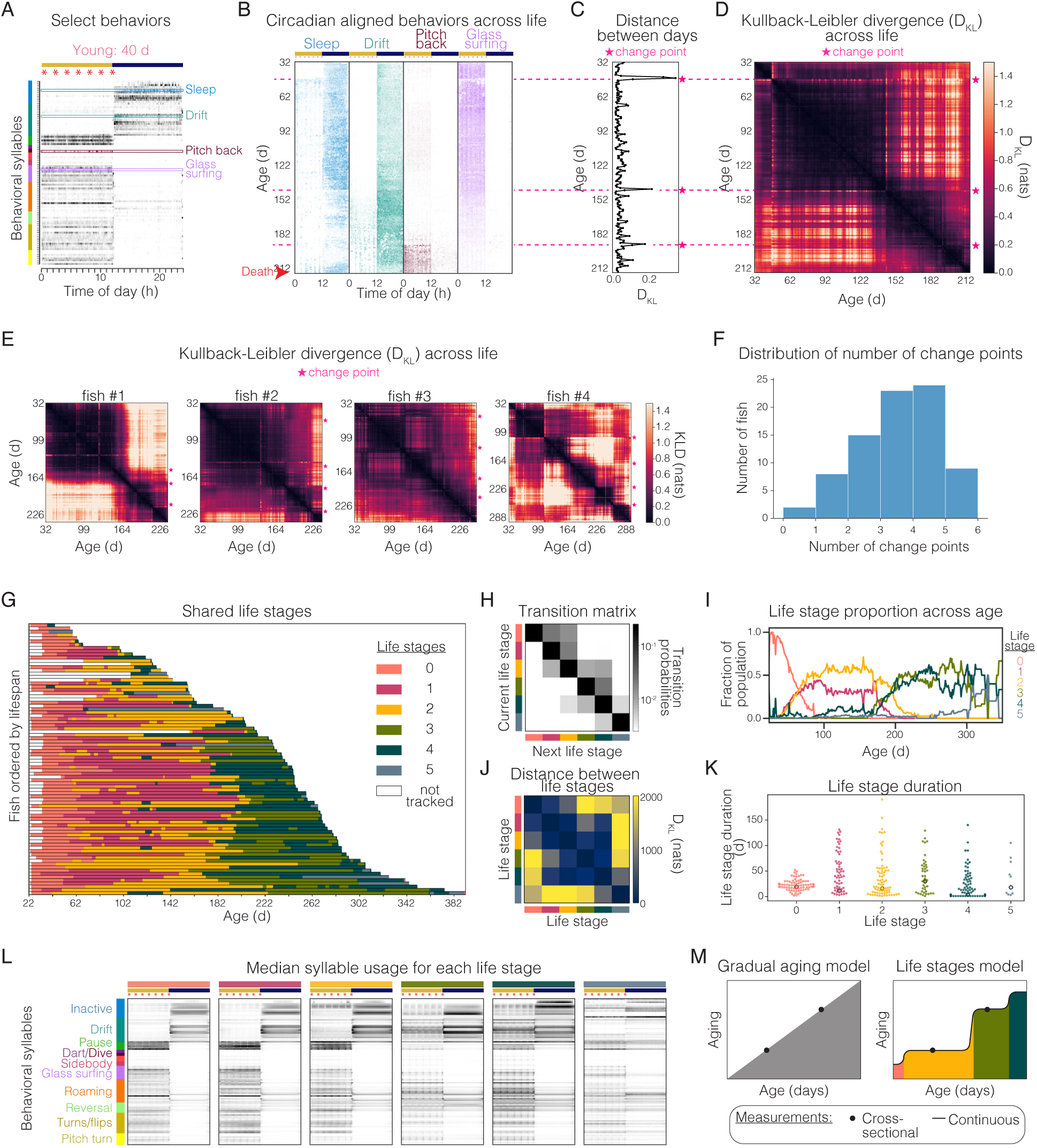
Aging animals exhibit stereotypical behavioral transitions across-life. (**A**) Single day (age 40 days) behavioral syllable use for a single animal highlighting four specific behavioral syllables. Throughout figure, ordering and clustered of behavioral syllables are as in Fig. 1I; red asterisks indicate feeding times; light/dark cycle is indicated by yellow/blue bar. (**B**) Circadian-aligned, whole-lifespan heatmaps of four select behaviors highlighted in **A**. (**C**) Behavioral distance (i.e., D_KL_) of behavioral syllable usage distribution comparing subsequent days highlighting detected change points (pink star and dashed line) which aligns with specific days with abrupt transition in behavioral syllable usage. (**D**) Cross-correlation using symmetrized D_KL_ of the behavioral syllable usage distribution comparing all days in one animal’s life to all other days ordered from youth (32 d) to death (231 d). Scale of days in **C** and **D** is aligned with age in **B**, and the same animal is shown in all. Change points are highlighted with a pink star. (**E**) Cross-correlation using symmetrized D_KL_ as shown in **C** for four animals ordered by lifespan. (**F**) Distribution of the number of change points observed per animal. (**G**) Survival curve of the population of tracked animals (n=81), with coloring of each animal based on HMM-derived life stage. (**H**) Transition probability matrix for each life stage. (**I**) Proportion of the population in each of the six life stages across age. (**J**) Symmetrized D_KL_ of HMM emission distributions across life stages. (**K**) Life stage duration; each point is a different animal. (**L**) Median 100-behavioral syllable use for animals in each life stage. (**M**) Comparison of models of aging.

To quantify these transitions, we performed cross-correlation analysis comparing each day in an animal’s life to all other days, and measured behavioral distance between days using symmetric Kullback–Leibler divergence (D_KL_) across all 100 behavioral syllables (Fig. 6C-D; fig. S18). We observed striking transitions at characteristic ages in each individual (Fig. 6B-E; fig. S18), while between abrupt transitions, we observed long periods of largely stable behavior composed of very similar days. To quantify these discrete transitions, we performed change point detection on behavioral syllable usage across life. For the animal depicted in Fig. 6A-D, we identified three discrete change points (highlighted in Fig. 6C-D) which align with peaks in D_KL_ from one day to the next (Fig. 6C-D). The first of these transitions occurred during relative youth (47 days), after which the animal shifted from exclusively sleeping during the night to exhibiting moderate day-time sleeping. The second transition occurred at more advanced age (148 days), after which the animal showed a reduction in time spent sleeping during the night. And the final transition occurred close to death (192 days), after which the animal exhibited increased time spent swimming pitching backwards (a likely maladaptive swimming behavior which could be due to changes in buoyancy or balance of unknown etiology (*72*)). Performing the same analysis for all animals across a cohort revealed abrupt transitions in behavior across individuals, with change points occurring at slightly different ages in different animals (Fig. 6E; fig. S18D). The distribution of discrete change points (transitions) detected across all animals reveals that the vast majority of animals go through at least one transition in their adult life while most go through three to four transitions (Fig. 6F). These observations suggested that animals progress through distinct, stable behavioral stages across the lifespan.

To investigate whether these stages are shared across animals, we modeled lifelong behavioral dynamics using a Gaussian HMM fit to the age factors. We identified six stable behavioral stages by cross validation (Fig. 6G; fig. S19A). However, we note that the precise number of stages may depend on the model specification. Intriguingly, these stages followed a natural order characterized by typically progressing forward, instead of switching back and forth between stages across days, illustrated by the transition matrix of the trained HMM (Fig. 6G-H). Given that these latent stages appear to be age-associated, we adopted the term *life stages* for long-duration stages that progress forward in the orderly fashion shown. Analysis of the proportion of the population in each life stage across age illustrates that very young animals start exclusively in one young-life stage (pink), afterward there is a split, with about half the animals entering one mid-life stage (yellow) and the other half entering an alternative mid-life stage (magenta). Likewise, there is a similar split into two old-life stages (dark green and light green) (Fig. 6I). Young-and mid-life stages were found to be more behaviorally similar to each other than to old-life stages, assessed by divergence (via D_KL_) between HMM emission distributions across life stages (Fig. 6J). Not all animals exhibited all six life stages, perhaps in part because life stages show variation in length across a population from a single day to >150 days (Fig. 6K).

To investigate the distinct behavioral properties of life stages, we evaluated the median daily behavioral syllable use of animals in each stage (Fig. 6L). We find that animals in young-(pink) and mid-life (yellow/magenta) stages exhibited inactive/sleep and drift behavior mostly at night, whereas animals in old-life stages (light and dark green) showed a shift in circadian regulation, with inactive/sleep and drift behavior persisting throughout both day and night (Fig. 6L). In late-life stages, there was a progressive reduction in pause behaviors in late-life. These pause behaviors typically indicate transitions from one behavior to another, suggesting that younger animals are more behaviorally flexible and dynamic (Fig. 6L). In the rare late-life stage (blue), animals exhibited a complete shift in behavior with a few specific behavioral syllables persisting constantly throughout the day and night (Fig. 6L). Recordings of animals in this rare late-life stage (blue) reveal an age-associated swim bladder disorder leading to disrupted buoyancy, which can be linked to infections in fish (*73*) (see Methods). This age-associated phenotype is observed sporadically in <10% of recorded animals (and is not enriched in any cohort). Thus, our unsupervised behavioral approach is able to identify animals in distinct, rare states that may be linked to disease.

We also noted an intriguing correlation between an animal’s transition into late-life stages (light and dark green) and a decline in feeding behavior measured either via feeding behavior syllables or by inspection of tanks for leftover uneaten food (fig. S19B). The drop in feeding is accompanied by a range of concurrent behavioral changes—loss of circadian regulation, increased inactivity, and drift behavior during both day and night (and likely represent poor health). These periods varied in length across individuals, but most animals entered into this state substantially prior to death. Longer-lived animals generally spent a larger proportion of life in this state, suggesting that healthspan and lifespan may be uncoupled at older ages in precisely quantifiable ways with this system, and that extending the healthy stages even at advanced ages (yellow and pink states) could be a well-defined and accessible quantitative target for extending human health. This life stage model (in which aging proceeds in discrete steps or life stages, in contrast to a gradual continuous decline; Fig. 6M) would be challenging to detect with cross-sectional measurements at different ages, supporting the importance of continuous and longitudinal study of aging.

## Discussion

Here we provide the first longitudinal, continuous, high-resolution, and scalable view of the process of aging in vertebrates through behavioral screening from adolescence until death. With this dataset, we observed that animals do not all age in the same way but, rather, follow distinct aging trajectories. The specific trajectory an individual takes through life, as well as the speed at which that path is traversed, is influenced by sex, longevity interventions, and future lifespan.

Intriguingly, the presence of distinct types of human aging (*74, 75*) supports generalizability of our observation in killifish. It may also be relevant that human beings exhibit lifelong remodeling of genome structure in neurons crucial for defining motor and behavioral patterns (*76*).

We discovered that even at a relatively young age, individuals destined for a short lifespan are behaviorally distinct from individuals destined for a long lifespan. For example, at a young age, animals destined for a long lifespan exhibit higher peak locomotor velocity (i.e., “sprinting”) than animals destined for a short lifespan. Interestingly, maximum velocity at middle age is highly correlated with future lifespan in *C. elegans* (*77*) and in humans, gait speed and decline in gait speed is associated with disability and mortality in older adults (*26, 78–80*). Together suggesting that there are characteristics of locomotion conserved across species that could be used early in life to predict future lifespan.

We also found that by middle age, there are strong molecular signatures linked to an individual’s aging trajectory and future lifespan. Specifically, transcriptomic analysis revealed elevated liver ribosome biogenesis in short-lived individuals. This result may be particularly interesting since less ribosome biogenesis is linked to longevity and longevity interventions (*81*) while enhanced ribosome biogenesis and translation are observed in fibroblasts from progeria patients (*82*).

Together, these data suggest that the level of ribosome biogenesis even early in life could be a key regulator of lifespan.

The striking observation that short-lived and long-lived animals traverse divergent aging trajectories indicates that even at an early age, one can distinguish short-lived and long-lived individuals solely on the basis of behavior. To test this hypothesis quantitatively, we trained classification models, which we found accurately predicted an animal’s future lifespan on the basis of behavior. Behavior is highly expressive of the aging process, in line with prior work in mice indicating that behavior and frailty metrics can be used to predict age and remaining life (*23, 83*), but given the longitudinal nature of our dataset, we were able (for the first time in a vertebrate) to show that at a young age (prior to mid-life), behavior alone can accurately forecast an animal’s future lifespan as short vs long. In future, it would be exciting to build models to accurately predict remaining lifespan as a continuous variable given behavior at a young age.

Analogous noninvasive recording in humans (via e.g., wearable devices (*84*)) might provide a window into long-term health and lifespan; for example, early and accurate prediction of lifespan and age-associated disease risk could help guide early intervention (*26, 84, 85*).

Our unique view into the aging process revealed a distinctive structure in which animals stably maintain one behavioral stage for a substantial fraction of the lifespan and then transition to another distinct (and similarly stable) behavioral stage. We observed little switching back and forth between stages, noting instead an orderly progression. These shared life stage patterns suggest an intriguing model of the aging process in which, over time, the effects of aging factors accumulate to the point that the animal must shift to a new distinct stage to reach a new stable homeostasis. This stage-based model of the architecture of aging resembles well-established models of embryonic development that include distinct staging (*1–3*), suggesting that perhaps such staging continues throughout life. This behavioral architecture could be driven in part by energy conservation decisions (*28*), which may be best implemented in a staged manner (analogous to a gear shift in having different discrete modes rather than continuously variable transmission).

While human beings exhibit the known discrete transitions of sexual maturity and menopause as well as sporadic health events throughout life (*86*), whether we can identify discrete stages of aging in human beings– analogous to those seen in killifish– remains unknown. However, supporting the notion of several life stages in humans, cross-sectional transcriptional profiling of the human frontal cortex across age revealed large changes in the fourth and eighth decades of life (*87*), and analysis of human plasma across lifespan revealed waves of changes in the proteome in the fourth, seventh and eighth decades of life (*88*), while multi-omic longitudinal profiling of human aging revealed nonlinear patterns in molecular markers of aging (with substantial dysregulation occurring at approximately the fourth and sixth decades of life) (*89*).

There is also evidence of non-linear changes to mitochondrial metabolism that may result in a staged structure of metabolic aging (*90*). These molecular shifts could be associated with transitions to distinct life stages (as with the genome structure changes noted above (*76*)), but this intriguing possibility remains to be established. Just as understanding and defining discrete staging has been fundamental to the mechanistic study of development (*1–3*), the staged model of aging architecture described here may provide a powerful approach to enable targeted mechanistic and therapeutic discovery work in the study of human aging.

## Material and Methods

### African turquoise killifish care and husbandry

African turquoise killifish *Nothobranchius furzeri* (GRZ strain) were maintained according to established guidelines (*45, 47, 91–93*). Briefly, killifish embryos, originating from a breeding tank with one male (1–3 months) and 4–5 females (1–3 months), were raised in Ringer’s solution (Millipore, 96724), with two tablets per liter of RO water and 0.01% methylene blue (i.e. embryo solution) at 26–27 °C in 60 mm x 15 mm petri dishes (Fisher Scientific, 07-000-328) at a density of <100 embryos per plate. After two weeks in embryo solution, embryos were transferred to moist autoclaved coconut fiber (Eco Earth Coconut Fiber, EE-8) lightly packed in petri dishes where they were incubated for another two weeks at 26–27°C. After 2–3 weeks on moist coconut fiber, embryos were hatched. For hatching, embryos were placed in humic acid solution (1 g/l, Sigma-Aldrich, 53680 in RO water) and incubated overnight at room temperature. Once hatched, animals were housed at 26–27 °C in a central filtration recirculating system (Aquaneering, San Diego, USA) at a conductivity between 1000–4000 μS/cm and a pH between 6.5–7.0, with a daily exchange of 10% water treated by reverse osmosis (i.e. RO water). Animals were kept on a 12-hr light/dark cycle. Young fish (i.e., pre sexual maturity) were fed freshly hatched brine shrimp (Brine Shrimp Direct, BSEP6LB) two times a day on weekdays and one time a day on weekends. Adult fish (i.e., post sexual maturity; ≥ 3–4 weeks of age) were fed dry fish food (Otohime fish diet, Reed Mariculture, Otohime C1), 5–7 mg per feeding, seven times a day for the *ad libitum* feeding paradigm or three times a day in the morning for the dietary restriction feeding paradigm using custom programable automated feeders (*50*). Fish were visually monitored daily for overall health. System water and tank detritus were submitted for PCR testing for aquatic pathogens. From these tests, throughout recordings, the system was negative for all *Mycobacterium* species tested with the exception of *M. chelonae* and *M. fortuitum* and also negative for *Pleistophora hyphessobryconis*, *Pseudocapillaria tomentosa*, *Pseudoloma neurophilia*, *Myxidium streisingeri*, and *Picornavirus*. During experiments including females and males under dietary restriction, bryozoans were present in the aquatic sump. They were manually removed weekly. We observed sporadic occurrence of an age-associated swim bladder disorder leading to disrupted buoyancy (commonly referred to as ‘belly sliders’). These animals were monitored daily to ensure they were able to swim to and consume food. There are different reports of the cause for this loss in swim bladder function. One proposed cause is due to mycobacterial infection derived from live food (e.g., hatched brine shrimp) (*73*). Other reports find that effected animals were negative for mycobacteriosis and instead found histopathology consistent with microsporidia and subsequent sequencing analysis revealed high levels of sequence homology with *Loma acerinae* (*72*). The length of lifespan of the African turquoise killifish has been reported to substantially vary between animal facilities (and within a facility over time), likely due to different husbandry conditions and other yet uncharacterized environmental parameters, which are common for a relatively recent model system (*45, 47, 91–93*). All animals were raised in accordance with protocols approved by the Stanford Administrative Panel on Laboratory Animal Care (protocol #APLAC-13645).

### System for life-long recordings

Animals were placed in standard 2.8-L tanks (Aquaneering, San Diego, USA) in a custom-built table rack system (Fig. 1A, fig. S1). Tanks received continuous water inflow and outflow from a central filtration recirculating system (Aquaneering, San Diego, USA) with chemical and biological filtration (with water parameters described above). Each table rack was built to hold twelve 2.8-L tanks with space for opaque acrylic dividers between tanks to prevent animals in adjacent tanks from seeing one another (fig. S1). At the center of the custom rack, an acrylic gutter (Tap Plastics, San Jose, USA) was fixed for system water outflow from tanks. The rack system was built so that the top of the tanks was 2 m from the bottom of the overhead mounted camera lens (fig. S1E). The distance of the camera from the tanks is important to prevent obstruction of the tank edges. One camera was used for each 12-tank table rack for a total of three cameras and three table racks (fig. S1A).

For recordings, overhead mounted infrared-sensitive Basal industrial cameras (aca2040, monochromatic, 2048×1536, Basler AG, Germany) with attached lenses (M0824-MPW2, Computar, USA) and infrared filters (Nr.2.3) were run continuously capturing at 20 frames per second (fps) run through the Motif Video Recording System (loopbio, Austria). Custom built infrared backlighting (LLPX-750X750-850-CSM, Smart Vision Lights, USA) was fixed under the tanks (fig. S1B). Infrared backlighting and filters were used to allow for recordings throughout the 12-hr day/12-hr night lighting cycle so that circadian changes in visible light would not influence the recording itself. Tanks were covered with a custom cut red acrylic cover (Chemcast transparent acrylic 1/8^th^ inch, Tap Plastics, San Jose, USA) to prevent animals from jumping out of tanks (fig. S1A-B). The red color acrylic did not obstruct the infrared light on its path to the camera. The cover is largely invisible in the recordings.

Male or female killifish were individually housed in recording tanks depending on the experiment starting once they reached sexual maturity as determined by the appearance of sexually dimorphic coloration (∼3–4 weeks of age; (*94*)). Animals lived in the same tank throughout the recording period (typically until death unless otherwise noted, for example, for molecular profiling) and recordings were collected continuously (20 fps) 23-hrs, 59-min and 30-s each day. Recordings were automatically stopped at a set time each day (11:21:43 am). The recordings were then reinitiated for the next 24-hrs of recording (11:22:13 am). The recorded frames from the completed day were then automatically transferred from the recording computer to a storage computer. Fish were enrolled in rolling cohorts to enable the largest number of animals to be tracked and to avoid empty tanks on the recording tables resulting in 10 cohorts (cohort 1=24 animals; cohort 2=15 animals; cohort 3=14 animals; cohort 4=8 animals; cohort 5=8 animals; cohort 6=18 animals; cohort 7=11 animals; cohort 8=8 animals; cohort 9=10 animals; cohort 10=3 animals; cohort 11=30 animals; and cohort 12=39 animals). The majority of fish were recorded until natural death however a small number of individuals were removed prior to natural death due to (1) jumping out of their tank, (2) removed to make space for a new cohort, or (3) removed for cross-sectional processing such as RNA sequencing. In total, 188 fish were tracked with 121 animals tracked until natural death, 30 females and 39 males under a dietary restriction feeding paradigm.

Fish and tanks were visually monitored daily for overall health and cleanliness. If tanks showed buildup of detritus, they were manually cleaned with Plastic Pasteur Pipettes (Global Scientific, USA). Throughout recordings, key environmental parameters e.g., room lighting, room temperature, water temperature, water salinity, and water pH were monitored and regulated. Water parameters were regulated through the central filtration recirculating system (Aquaneering, San Diego, USA).

### Key point tracking

To track animals throughout recordings we used Loopy Video Analysis and Tracking software (loopbio, Austria). Specifically, six key points (snout, midbody, endbody, tail, fan, and sidebody) along the killifish body were manually annotated in frames of video recording. Manual annotations were performed on fish of diverse age and cohort and at different times of day to ensure the training annotations would generalize well for the whole dataset.

11,100 frames of individual fish were manually annotated in total. Not all key points were visible in each frame depending on the animal’s body positioning so the number of total annotations for each key point varied depending on frequency of key point visibility. For example, the number of annotations for each key point are as follows: 11,029 snout, 10,716 midbody, 10,609 endbody, 11,064 tail, 11,074 fan, and 466 sidebody. Manual annotations were used to train a deep convolutional neural network to predict key point positions for fish in unseen frames using Loopy Video Analysis and Tracking software (loopbio, Austria). Trained models were used to predict key points in all frames of the recordings using Loopy Video Analysis and Tracking software (loopbio, Austria). For each frame of recording, the Loopy Video Analysis and Tracking software (loopbio, Austria) would provide the x-and y-coordinates for each visible key point as well as a corresponding timestamp for the frame. The same key point tracking model was used for predictions across all recordings.

Each recording camera included up to 12 distinct animals because there are 12 tanks within the field of view of a single camera. All animals within the field of view were tracked simultaneously using Loopy Video Analysis and Tracking software (loopbio, Austria).

Individual animal key points were separated based on the coordinates of each tank’s edge. For individual tanks, if duplicate key points were detected (e.g., two snouts) due to a tracking mistake, that frame of the recording was dropped for that animal. Key point coordinates were shifted such that the origin was the bottom left corner of each tank and key point coordinates were rotated so that all tanks were oriented in the same direction (i.e., feeding side was the same for all fish).

### Pose features

Pose features were calculated from the x-/y-coordinates of tracked key points as well as the timestamp information from each frame of the recordings.

***Coordinates*** (‘x_snout’, ‘y_snout’, ‘x_midbody’, ‘y_midbody’, ‘x_endbody’, ‘y_endbody’, ‘x_tail’, ‘y_tail’, ‘x_fan’, ‘y_fan’). Raw x-/y-coordinates for each key point were shifted such that the origin was the bottom left corner of each tank and key point coordinates were rotated so that all tanks were oriented in the same direction (i.e., feeding side was the same for all fish).

***Velocity*** (‘snout_velocity’, ‘midbody_velocity’, ‘endbody_velocity’, ‘tail_velocity’, ‘fan_velocity’). Velocity was calculated for all key points after first filtering data using scipy.signal.lfilter to alleviate the impact of key point jitter.

***Acceleration*** (‘snout_acceleration’). Acceleration was calculated for the snout key point after first filtering data using scipy.signal.lfilter to alleviate the impact of key point jitter.

***Dispersion*** (‘disp’). The dispersion area is the area of the smallest bounding circle enclosing the snout key point within a defined time window. Our set time window length is two seconds (i.e., 40 frames). Dispersion was calculated using the max and min x-and y-coordinate of the snout within the select time window.

***Bounding area*** (‘bounding area’). The bounding area is the smallest bounding circle enclosing all visible key points of the whole fish in a defined time window. Our set time window length is two seconds (i.e., 40 frames). The bounding area is then centered at zero by subtracting the mean and normalized by the mean bounding area for that day.

***Normalized body length*** (‘body_length_norm’). Body length was calculated by summing the length of each body segment snout to midbody, midbody to endbody, endbody to tail, and tail to fan to get full body length. To calculate normalized body length, the body length was then centered at zero by subtracting the mean and normalized by the mean body length for that day.

***Body segment proportions*** (‘tail_fan_prop’). Measure of the proportion of an animal’s body length from the tail-fan segment.

***Key point count*** (‘count_snout’, ‘count_midbody’, ‘count_sidebody’, ‘count_endbody’, ‘count_tail’, and ‘count_fan’). Binary count of whether each key point is visible or not in a given frame (i.e., one if visible, zero if not visible). This was done for all key points.

***Snout distance from tank edge*** (‘dist_snout’). Positioning of the fish in the center vs. tank edge is important in differentiating several key behaviors (e.g., glass surfing behavior). Thus, we calculated the distance of the snout from the nearest tank edge.

***Heading direction*** (‘heading_direction’). The direction of the vector from midbody to snout.

***Body connectivity*** (‘alpha1’, ‘alpha2’, ‘alpha3’, ‘alpha_all’). The angle between body segments. For example, the angle between snout-midbody-endbody (alpha1), the angle between midbody-endbody-tail (alpha2), and the angle between the endbody-tail-fan (alpha3) body segments.

***Body curvature*** (‘theta1’, ‘theta2’, ‘theta3’). The local tangent direction at each key point – midbody (theta1), endbody (theta2), and tail (theta3) relative to heading. Tangent angles are positive when pointing toward the animal’s left side and negative when pointing toward the animal’s right side.

***Reversal*** (‘dot_product’, ‘reversal’, ‘reveral_binary’). To assess if the animal is moving forward or in reverse, we calculated the dot product of the heading direction vector and the snout velocity vector. For this, smoothing of the key-point coordinates was done first by rolling average over a 10-frame window.

***Sleep*** (‘sleep’). Sleep is behaviorally defined in zebrafish as periods of inactivity lasting one minute or longer (*95, 96*). We defined inactivity based on dispersion. If the dispersion area was similar to the movement expected from key point jitter over the course of two seconds (threshold for dispersion area is 50), then the animals was determined to be inactive. If the animal maintained this inactive state for greater than one minute, it was considered sleeping.

The above features were calculated across all frames. Then, depending on the feature, the mean and/or standard deviation of each was evaluated over a rolling window of 10 frames (Table S1) resulting in 57-dimensional vector per frame describing all pose features. To quantify whether distinct pose features were used during day/night or across different ages (Fig. 1G) for select animals, we measured linear separability as classification accuracy of a linear kernel support vector classifier fitted on the top five pose feature principal components (PCs) using sklearn.svm.SVC() with kernel=‘linear’ (*97*). For pose feature principal component analysis (PCA), we standardized (i.e., z-scored; subtracted mean and divided by standard deviation) the pose features prior to PCA (sklearn.decomposition.IncrementalPCA).

### Hidden Markov model (HMM) for behavioral syllables

We infer stereotyped, reused behavioral syllables from pose features by fitting a hidden Markov model (HMM) to a dimensionality-reduced representation of the pose features. The unprecedented amount of data (>30 billion frames) made it prohibitive from both a memory and computational time perspective to apply standard algorithms for fitting to the full dataset. The data, however, is highly autocorrelated across days and for a given individual. This supports the more data efficient approach of building training and validation sets on representative subsets of the data. The training set was built from a random selection of 200 full days of recording from different individuals, different cohorts, and at different ages (3.46×10^8^ frames at 20 fps resolution). This training set size still required incremental, distributed, and stochastic variations of algorithms, as described below. The validation set consists of recordings drawn from across ages but from individuals not present in the training set (2.1×10^7^ frames at 20 fps resolution).

Given the redundancy of many of the pose features and to improve the efficiency of HMM model training, we first standardized (i.e., z-scored; subtracted mean and divided by standard deviation) the pose features and then performed principal component (PC) analysis (PCA) (sklearn.decomposition.IncrementalPCA). The top 15 PCs were found to explain >80% variance (fig. S2A) and the standardized pose features were reduced down to these principal components. The resulting features were smoothed with a 10-frame moving average filter.

Finally, a Gaussian hidden Markov model (HMM) was fit to these preprocessed PCs to infer the underlying discrete hidden states, which are interpreted as “behavioral syllables.” The Baum-Welch algorithm (*98*) is the standard expectation-maximization (EM) algorithm for estimating the maximum likelihood of parameters in HMMs, but each iteration requires memory that is linear in the number of frames and quadratic in the number of states, i.e. *O*(*TK* + *K*), for *T* frames and *K* hidden states (>138 GB per state for each training iteration for 200 full days of recording). Note that in regimes like ours, in which *T* ≫ *K*, this simplifies to *O*(*TK*). To reduce the memory footprint of the standard implementation, which is prohibitive even for the reduced size of the training dataset, we developed a custom JAX implementation of stochastic EM for hidden Markov models (*99, 100*). This reduces the memory footprint to *O*(*BK*^2^), for batch size of) sequential frames, where *B* ≪ *T* and is adjustable based on machine constraints. This implementation also supports parallelization across batches, reducing the time complexity linearly in the number of parallel jobs. At each iteration, we run the forward-backward algorithm to infer the hidden states for a batch of size *B* using the current model parameters. Then, we perform a Robbins-Munro style update of the parameters by taking a convex combination of the previous parameters and the optimal parameters for the hidden states of the current batch. Our implementation using 71 batches required 2 days for computation using 16 GB/CPU and 8 CPU on a single node to fit an HMM with 100 hidden states. The custom implementation is available at https://github.com/lindermanlab/scalable-gaussian-hmm.

The primary hyperparameter in this model is the number of HMM states. We selected this hyperparameter by cross-validating against the log likelihood of the held-out validation dataset under the fitted model. Held-out log likelihood was observed to begin plateauing at 100 states (fig. S4A); model training was prohibitively slow for greater than 100 hidden states, so 100 hidden states were used for analysis. Behavioral syllables were predicted for the whole dataset using the fitted HMM for downstream analysis. The distribution of syllable duration varied depending on the syllable. We evaluated mean, 95^th^ percentile, and 5^th^ percentile syllable duration for all 100 syllables on single days comparing young (45 d) vs old (270 d) (fig. S4G) based on nine animals (five young and four old) presented in Fig. 1G.

### Uniform Manifold Approximation and Projection for Dimension Reduction (UMAP) embeddings to visualize pose features

UMAP embeddings of the top 15 PCs describing killifish pose features were used for visualization. Embeddings were built for select animals on a select day to build visualizations in Fig. 1G-H, fig. S3, and fig S4C-E. Due to the large number of frames, embeddings were built using 1 in every 5 frames (i.e., down-sampling from 20 Hz to 4 Hz frame rate) of the select data to avoid memory limitations. Embeddings were built using the top 15 PCs as described above (not smoothed) with the umap python package with n_neighbors=15, min_dist=0.3, n_components=2, random_state=42, metric=’euclidean’.

### Kullback-Leibler Divergence (D_KL_) analysis

Kullback-Leibler Divergence (D_KL_) analysis (relative entropy) was performed to calculate the pairwise dissimilarity. We use D_KL_ in three applications in the paper: (1) measuring dissimilarity of HMM-defined behavioral syllable emission distributions to establish behavioral syllable clusters (fig. S4B), (2) measuring dissimilarity of behavioral syllable usage distributions from each day in an animal’s life compared to all other days of the same animal’s life (Fig. 6C-E), and (3) measuring dissimilarity of HMM-defined life stage emission distributions (Fig. 6J).

For (1), first HMM-defined behavioral syllable emission distributions were built from the behavioral syllable emission mean and covariance using torch.distributions.MultivariateNormal. The pairwise KL divergence across all 100 behavioral syllables was calculated using torch.distributions.kl.kl_divergence. We then calculated symmetrized 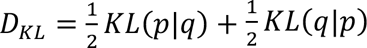.

For (2), to calculate the daily behavioral syllable usage distribution for a given day and for a given animal, we summed the total time spent in each of the 100 behavior syllables during the 12-hr light period and the 12-hr dark period to generate a 200-element long vector. This vector describes the daily behavioral syllable usage distribution for a given day in each animal’s life. D_KL_ between each day in an animal’s life was then calculated for all possible pairwise combinations of days within a single animal using the scipy.stats.entropy module. We then calculated symmetrized D_KL_ as described above.

For (3), HMM-defined life stage emission distributions were built from the life stage emission mean and covariance using torch.distributions.MultivariateNormal. The pairwise KL divergence across all six life stages was calculated using torch.distributions.kl.kl_divergence. We then calculated symmetrized D_KL_ as described above.

### Behavioral syllable clustering

To aid in interpretation and visualization of behavioral syllables, we perform hierarchical clustering of the behavioral syllable emission distributions. Hierarchical clustering was performed using sklearn.cluster.AgglomerativeClustering with distance calculated using symmetrized D_KL_ between syllable emission distributions (described above). The resulting clusters are visualized in fig. S4B.

### Tensor component analysis (TCA)

For dimensionality reduction of daily behavioral data, we used tensor component analysis (TCA), which is closely related to canonical polyadic (CP) decomposition (*101*), following methods and software described in detail in Williams A.H., et al. (*55*) (https://github.com/neurostatslab/tensortools/). We first binned the data at 10 min resolution by counting the occurrence of each syllable over 12000 frames (10 min). We chose to use 10-min binning because at this resolution we were able to capture key circadian events such as each feeding and the abrupt response to the light change while still reducing the size of the data for downstream processing. Nonnegative TCA models were trained using the 100 behavioral syllables binned at 10 min resolution for all animals and all days of recording. Usage traces for each behavioral syllable were normalized to range between zero and one across the whole dataset to avoid bias due to highly used behavioral syllables. Data was organized into an *N* × *T* × *K* tensor with 1 = 100 behavioral syllables recorded at *T* = 144 time bins/day over *K* = # animals × # days of recording/animal. We then fit nonnegative tensor decomposition models using normalized behavioral data. To select the optimal number of components, we performed cross-validation by holding out 50% of time-bins from the dataset and varying the number of tensor components from 2 to 100. As described by Williams *et al.*, we normalize the reconstruction error between zero and one to provide a metric analogous to the fraction of unexplained variance often used in PCA (Fig. 2B) (*55*). To evaluate models, the normalized reconstruction error was calculated for both the training and the held-out set. We selected a 45-component model for downstream analysis.

TCA approximates the data as a sum of outer products of three vectors—in our case: a) time-of-day factors, b) behavioral-syllable factors and c) age factors (Fig. 2A, D). Time-of-day and behavioral-syllable factors constitute structure that is common across all days in the dataset.

What makes one day different from another for each animal is the amplitude or weighting of each time-of-day and behavioral-syllable factor, i.e., the age factor. The age factor can be thought of as the amplitude/weight for a given animal at a given age in the daily behavior pattern. Some TCs ramp up or down during the day and night (e.g., TC19, TC22, TC41), whereas other TCs show constant weighting throughout the day and night, with no circadian regulation (e.g., TC24) (Fig. 2D, subpanel a). Behavioral-syllable factors vary from sparse (i.e., few syllables dominating, e.g., TC34) to broad (i.e., with many behavioral syllables making small contributions to build the component, e.g., TC43) (Fig. 2D, subpanel b). One tensor component (TC20) peaks at each of the seven feedings and is composed of behavioral syllables that represent feeding behavior (although additional direct measures of food consumption would be necessary to explore feeding in a more finely resolved manner). Two TCA models were trained, one model including tracking data from all males under the *ad libitum* feeding paradigm and a second model additionally including tracking data from females and males under dietary restriction. Age factors for the first model (including only males under the *ad libitum* feeding) are presented in Fig. 2, Fig. 3, Fig. 4 and Fig 6 while the second model age factor weightings for all animals are presented in Fig. 5.

### Aging trajectories

To visualize aging trajectories (i.e., the change in age factor weighting across life) for each animal, we performed PCA of TCA-derived age factors. Prior to PCA, we z-scored the TCA-derived age factors. We visualized the top three PCs (∼40% explained variance; fig. S7A) in Fig. 2E-G, Fig. 3B, Fig. 3H, Fig. 5D-E, Fig. 5G-H, fig. S7B, fig. S8, fig. S9, fig S11B-C, and fig. S16G. For visualization smoothed trajectories, we used scipy.ndimage.gaussian_filter. For visualization purposes only, we also performed UMAP embeddings of TCA-derived age factors. To build UMAP embeddings, we used the top 20 PCs (described above) using the umap python package with n_neighbors=15, min_dist=0.3, n_components=2, random_state=42, metric=’euclidean’. The resulting visualizations are shown in fig. S10. To evaluate how similar behavioral aging trajectories are across individuals, we quantified the summed variance of all 45 TCs at each age across life (fig. S10A).

### Organ harvesting

Animals were fasted for 24 hours before organs were harvested. RNA-sequencing was performed on 8 organs (brain, heart, intestine, kidney, liver, skin, spleen, and testis) in wildtype animals at different ages: young (80 d; n=8), middle-aged (150 d; n=17), and old (210 d, n=4). Each cohort was tracked starting at adolescence until the designated endpoint. Organs were harvested in the morning. Organs were dissected on ice-cold Sylgard-coated Petri dishes (filled with ice and covered in plastic wrap) and were snap-frozen in liquid nitrogen after harvesting (stored at-80°C until RNA isolation). Skin samples were collected from the caudal fin.

### RNA isolation

RNA was isolated from organs using QIAGEN RNeasy Mini kit (QIAGEN, 74106), following protocols previously described (*50*). Briefly, organs were transferred to 1.2 mL Collection Microtubes (QIAGEN, 19560), on dry ice to reduce tissue thawing. Autoclaved metal beads (QIAGEN, 69997) and 700 µL of QIAzol (QIAGEN, 79306) were added to each tube followed by tissue homogenization on a TissueLyserII machine (QIAGEN, 85300) at 25 Hz, 5 min each, at room temperature. Lysates were transferred to 1.5 mL tubes and 140 µL chloroform (Fisher Scientific, C298-500) was added followed by vortexing and incubation at room temperature for 2–3 min. Lysates were then centrifuged at 12,000×g at 4°C for 15 min. The aqueous phase was mixed with 350 µL ethanol (200 Proof, Gold Shield Distributors, 412804) and then transferred to RNeasy columns from the RNeasy RNA Purification Kit (QIAGEN, 74106). The RNeasy column was then washed with 350 µL RW1 buffer (provided by the RNeasy Mini kit), and treated with DNase I (following the kit’s protocol) at room temperature for 15 min. The column was washed two times with 500 µL RPE buffers, and the RNA was eluted with 50 µL nuclease-free water (Invitrogen, 10977023). RNA quality and concentration were measured using an Agilent 2100 Bioanalyzer and the Agilent Nano Eukaryotic RNA Kit (Agilent, 5067-1511). All bioanalyzer assays were performed by the Stanford Protein and Nucleic Acid Facility.

### Multi-organ RNA-sequencing analysis

***cDNA library generation and sequencing using BRB-seq for RNA barcoding*.** Organ RNA samples were prepared following manufacture recommendations for the BRB-seq platform with the Mercurius™ Protocol (Alithea Genomics) for bulk RNA barcoding and sequencing. Briefly, RNA concentration was determined by using the Qubit 1X dsDNA High Sensitivity Assay Kit (Thermo Fisher, 797 Q33231) and between 0.48 and 0.66 µg of total RNA from each sample were reverse transcribed in a 384-well plate with unique barcoded oligo-dT primers. Amount of loaded RNA was matched between all samples of a given tissue but amount varied between different tissues based on yield from RNA isolation (brain-0.66 µg; gut-0.66 µg; heart-0.48 µg; kidney-0.66 µg; liver-0.66 µg; skin-0.48 µg; spleen-0.48 µg; testis-0.66 µg per sample). For sample pooling, 10 µL of each barcoded reverse transcribed sample were pooled and purified using Zymo Clean & Concentration Kit (Zymo, D4014) column and DNA Binding buffer (Zymo, D4004-1-L) followed by free primer digestion. Double-stranded cDNA was generated by second-strand synthesis via the nick translation method. The concentration was measured by Qubit (190 ng/µL), and prepared for tagmentation using the Tn5 transposase pre-loaded with adapters for library amplification. The cDNA library was prepared for Illumina sequencing and sequenced on an Illumina NovaSeq 6000 (2×150 bp paired-end) by Novogene (Novogene, Beijing, China), at a sequencing depth of >16 million pair-end reads per sample.

***RNA-seq data processing pipeline*.** Adaptors were first trimmed from raw sequencing FastQ files using Cutadapt (version 3.1) for removing the last 122 bases of each read (with parameters `-u-122`) followed by quality assessment using FastQC. Processed reads were then aligned to the African turquoise killifish reference genome (Nfu_20140520, GCF_001465895.1 (*39*)) and the gene read count and UMI count matrices were created using STAR (version 2.7.1a) with parameters adjusted according to recommendations for the BRB-seq platform with the Mercurius™ Protocol (Alithea Genomics) including:

*--soloCBwhitelist*: text file with list of barcodes used by STAR for demultiplexing.

*--soloCBstart*: start position of the barcode in the R1 fastq file = 1.

*`--soloCBlen`*: length of barcode = 14

*`--soloUMIstart`*: start position of UMI = 15

*`--soloUMIlen`*: length of UMI = 14

The sequenced library had 86.3% of reads mapped uniquely to the genome. Raw gene expression values were then normalized using DEseq2 (version 1.36.0) excluding genes with < 20 counts across all samples. For differential expression analysis between young (80 d) and old (≥210 d) animals, we used DEseq2 with the design ‘∼ age’ and then evaluated the differences between young (80 d) and old (≥210 d) animals. For differential expression analysis between predicted short-lived and long-lived behavioral trajectories, we used DEseq2 with the design ‘∼ trajectory’ for only the middle aged (150 d) data and then evaluated the differences between predicted short-vs. long-lived behavioral trajectory animals (Table S2). PCA was performed after filtering out genes with < 20 counts across all samples followed by variance stabilizing transformation (vst function in the DEseq2 package) (Fig. 3I, fig. S13A). The resulting filtering led to the following gene numbers per tissue: 18,723 (liver), 20,773 (brain), 19,335 (intestine), 19,158 (heart), 19,995 (kidney), 18,725 (skin), 20,070 (spleen), and 21,789 (testis).

***Enrichment analysis.*** To perform over-representation analysis for significantly differentially expressed genes between either young (80 d) vs. old (≥210 d) or short-lived vs. long-lived trajectory (all 150 d old), we used Gene Ontology (GO) enrichment analysis using the enrichGO function in the clusterProfiler package (version 4.4.4) (Table S2). For young vs. old, we ran analysis on genes significantly upregulated (padj<0.05 & log2FoldChange>0) and significantly down regulated (padj<0.05 & log2FoldChange<0). For short-lived vs. long-lived trajectory, we ran analysis on genes significantly upregulated (pvalue<0.05 & log2FoldChange>0) and significantly down regulated (pvalue<0.05 & log2FoldChange<0). For gene set enrichment analysis, we used GO analysis using the gseGO function in the clusterProfiler package (version 4.4.4) (Table S2). Genes were ranked by sign(log2FoldChange)*(-log10(padj)). Each killifish gene was then assigned to their human ortholog (best hit protein with BLASTp E-value>1e-3). Killifish genes with no human ortholog were removed. Where multiple killifish paralogs were present for a single human ortholog, the killifish paralog with the smallest adjusted p-value was kept. Enrichment analysis was run using the Bioconductor annotation data package (org.Hs.eg.db v3.15.0). The p-values of the enriched pathways were corrected for multiple hypotheses testing using the Benjamini–Hochberg method.

***Clustering/separability of transcriptomes of short-vs long-lifespan trajectory animals.*** To evaluate separation of transcriptomes of the short-vs long-lifespan trajectory animals across all tissues, we measured silhouette score (sklearn.metrics.silhouette_score) in PC1/PC2 space of true labels (long-lifespan vs short-lifespan trajectory based on behavior) compared to a null distribution built by permuting randomized group assignment across all n=17 samples to two groups and evaluating the resulting silhouette score (fig. S13B). We also performed unbiased k-means clustering of the liver PC1/PC2 data using sklearn.cluster.KMeans with the number of clusters set to two and default parameters which resulted in two clusters that matched the observed short-lifespan and long-lifespan trajectory labels (fig. S13C).

### Regression models (i.e., behavioral clocks) to predict age

Our goal was to build models to accurately predict animal age given the behavioral characteristics of the animal on a given day. Models were designed so that each day is described by a vector of the 45 TCA-derived age factors used as input features (no smoothing or standardization was performed) and then predict the animal’s age. We performed leave-one-fish-out cross validation where one fish in the training set was held-out, the model was trained on all days for remaining fish using input features and age label, and then predictions were made on the held-out fish based on input features (Fig. 4B-C). For cross validation, random forest regressors were built using sklearn.ensemble.RandomForestRegressor with default parameters. To evaluate model performance, we evaluated correlation of model predicted age with true age for all ages of held-out fish using Pearson correlation with numpy.corrcoef (Fig. 4B). We evaluated the absolute value of the error of held-out animal age predictions vs. true age for each age tested and then calculated the median absolute error across all animals for each age (Fig. 4C).

We also trained random forest regressors and evaluated model performance on a held-out test set of animals. For model training, random forest regressors were built using sklearn.ensemble.RandomForestRegressor with n_estimators = 500 and default parameters.

Held-out test-set animals came from three separate cohorts run at different times on the tracking system including – cohort 5 (8 animals tracked for their whole lifespans), cohort 6 (18 animals tracked until 4 months old) and cohort 9 (10 animals tracked for their whole lifespans) for a total of 36 held-out animals. No animals from any of these cohorts were used for model training.

Model performance was then evaluated by correlation (fig. S15B) and median absolute error (fig. S15C) as described above. To investigate which of the 45 TCA-derived age factors were most important for model predictions, we measured the mean decrease in impurity using the sklearn.ensemble.RandomForestRegressor attribute feature_importances_ on the trained regressor (Fig. 4D).

To predict age in cohorts of males under dietary restriction and female, because these datasets were analyzed using a distinct TCA model, we trained a new random forest regressors trained as described above on data from the second TCA model (using sklearn.ensemble.RandomForestRegressor with n_estimators = 500 and default parameters). For training, we held-out a test set of males animals under *ad libitum* feeding described above (cohort 5, n=8 animals tracked for their whole lifespans; cohort 6, n=18 animals tracked until 4 months old; cohort 9, n=10 animals tracked for their whole lifespans) as well as all males exposed to dietary restriction (n=39 animals tracked until 4 months old) and all females (n=30 animals tracked for their whole lifespan) such that the model was trained exclusively with data from males under *ad libitum* feeding. Model performance was then evaluated on the held-out test set of males under *ad libitum* feeding by both Pearson correlation (R = 0.93) and median absolute error across all ages (14 days). This model was then used to predict age of males under dietary restriction (Fig. 5F) and females under *ad libitum* feeding (Fig. 5I).

To compare the rate of aging of individual animals (fig. S16) we calculated the best fit linear slope of behavior clock predicted age vs true age (using scipy.stats.linregress) for each individual across all days recorded. We then compare the DR fed animals to the *ad libitum* fed animals. For the *ad libitum* fed animals we used a held out test set of animals not used for behavior clock training. DR fed animals had a significantly slower rate of aging compared to the *ad libitum* fed animals (P<0.0001 Mann Whitney test; fig. S16H).

### Classification model to forecast future lifespan

Our goal was to build models to accurately forecast an animal’s future lifespan as either long-lived or short-lived based on their behavior. To do this, we built separate classification models at ages throughout the killifish lifespan. We chose to build age-fixed classification models to detangle the impact of age on prediction accuracy given that age alone is a relatively good predictor of remaining life. To avoid day-to-day noise in behavior, we built each age-specific model taking the mean behavior over a set window of days (a 5-day window). Thus, each age-specific model was designed to use the mean of the 45 TCA-derived age factors over a 5-day window prior to the prediction age as input features and then classify the animal as either short lived or long lived. Input features were standardized (i.e., z-scored; subtracted training set mean and divided by training set standard deviation). We defined animals as being long lived if they live equal to or longer than 200 days (33^rd^ percentile). We selected 200 days as the cutoff as this represented a significant deviation from the median lifespan of the population (243 days) but a large enough fraction of the population for model training. We excluded extreme agers—animals that lived for >300 days (90^th^ percentile). For age-specific classifiers, we used random forest classifiers with sklearn.ensemble.RandomForestClassifier with default parameters. Each age-specific model was trained using leave-one-fish-out cross validation (as described above).

Leave-one-fish-out model performance was evaluated by prediction accuracy (using sklearn.metrics.accuracy_score) and Area Under the Receiver Operating Characteristic Curve (ROC AUC using sklearn.metrics.roc_auc_score) for each age-specific model (Fig. 4F-G). To evaluate the level of separation between the model predicted short-lived vs. long-lived groups, we grouped animals by held-out predictions at a set prediction age and then plotted the measured Kaplan-Meier survival curves of these two populations of animals using lifelines.KaplanMeierFitter (Fig. 4H). This was done for three select prediction ages: 70, 90, and 110 days. We then evaluated whether the predicted short-lived and predicted long-lived groups have a significant separation in true lifespan using a Log-rank test (lifelines.statistics.logrank_test).

To better understand the behavioral differences in animals destined for a short lifespan vs. long lifespan, we performed differential behavioral analysis between short-lived and long-lived animals across ages from 50-150 days old. For this analysis, we measured the mean behavior of short-lived animals and long-lived animals (i.e., the 45 TCA-derived age factors) across age. Significance was determined using Mann–Whitney U test with Bonferroni correction (with scipy.stats.mannwhitneyu and statsmodels.stats.multitest.multipletests) (Fig. 4I).

To classify lifespan in females, because these datasets were analyzed using a distinct TCA model, we trained a new age-specific (70-days old) random forest classifier trained as described above on data from the second TCA model using sklearn.ensemble.RandomForestClassifier with default parameters. Model performance was evaluated by leave-one-fish-out predictions based exclusively on behavior of males under *ad libitum* feeding by both prediction accuracy=0.77 (using sklearn.metrics.accuracy_score) and ROC AUC=0.72 (using sklearn.metrics.roc_auc_score). This model was then used to classify lifespan of females based on behavior at 70 days old (fig. S17F).

### Change point detection

To evaluate the number of discrete transitions in behavior across life, we used change point detection analysis to identify discrete times within individual aging trajectories in which there was a shift in underlying characteristics of the time series. We performed change point detection on daily behavioral syllable usage across life. To calculate the daily behavioral syllable usage distribution for a given day and for a given animal, we summed the total time spent in each of the 100 behavior syllables during the 12-hr light period and the 12-hr dark period to generate a 200-element long vector. This vector describes the daily behavioral syllable usage distribution for a given day in each animal’s life. We smoothed daily behavioral syllable usage to alleviate the impact of daily noise using scipy.ndimage.gaussian_filter with sigma=1 and then standardized the smoothed age factors (i.e., z-scored; subtracted mean and divided by standard deviation). We performed change point analysis using the python package rupture; https://github.com/deepcharles/ruptures (*102*). Kernel change point detection (*103, 104*) was performed with a gaussian kernel (using rupture.KernelCPD), an unknown number of change points per animal, and a penalty that identically scales with the natural log of lifespan length across all animals. We indicated change points in individual example animals in Fig. 6C-E. We evaluated the number of discrete change points observed across the life of each animal and the plot of the distribution of change point number per animal across the whole population is shown in Fig. 6F.

### Hidden Markov model (HMM) for life stages

To build the life-stage model, we first smoothed the TCA-derived age factors to alleviate the impact of daily noise using scipy.ndimage.gaussian_filter with sigma=1 and then standardized the smoothed age factors (i.e., z-scored; subtracted mean and divided by standard deviation). We trained Gaussian HMMs with *k*-means initialization and iterated until convergence (n=20 iterations of expectation–maximization) using Dynamax (a library for probabilistic state space models written in JAX: https://github.com/probml/dynamax). We varied the number of HMM states and observed the held-out log probability plateaued around 5 states and dropped off sharply at 7 states (fig. S19A). The resulting life stage model with six states accurately discriminated a rare life stage when animals are impacted by a swim bladder disorder leading to disrupted buoyancy which was missed in the five-state model. Given the biological relevance of this rare sixth state, we chose to move forward with a six-state HMM for life stage modeling. Using full covariance vs. diagonal covariance gaussian HMMs could yield a different number of stages.

We next investigated the characteristics of this six-state life-stage model. To evaluate the empirical transition matrix of the trained HMM, we counted the frequency of each state-to-state transition across all animals in the cohort (Fig. 6H). To evaluate the length of each life stage, we calculated the empirical duration of each bout in each life stage across all animals and evaluated median duration in each life stage (Fig. 6K). And to investigate the behavioral syllable signatures of each life stage, we grouped days by life stage and evaluated median time in each behavioral syllable across all time bins throughout the 24-hr day for each life stage. This was done with 10-min binned behavioral syllable data (i.e., 144-time bins per day) (Fig. 6L).

## Acknowledgements

We thank Drs. Felix Boos, Daniel Heinzer, and Daniel Richard as well as Angela Pogson, and all members of the Brunet lab and Deisseroth lab for input on the project and feedback on the manuscript. We thank Rogelio Barajas, Jacob Chung, and Sarah Boyle for excellent killifish husbandry support. We thank Emily Nguyen and Josh Moon for support setting up and checking on tracking cohorts. We thank Dr. John Bedbrook for design and construction of the custom racks for housing tanks during recordings. We thank Rahul Nagvekar for checking on fish in the tracking systems at various points in the project. This work was supported by R01AG063418 (A.B. and K.D.), a Knight Initiative for Brain Resilience Catalyst Award (A.B., S.W.L. K.D.), the Keck Foundation (K.D.), the ARIA Foundation (K.D.), the Glenn Foundation for Medical Research (A.B.), the Simons Foundation (A.B.), Chan Zuckerberg Biohub – San Francisco (A.B.), a NOMIS Distinguished Scientist and Scholar Award (A.B.), the Helen Hay Whitney Postdoctoral Fellowship (C.N.B.), the Wu Tsai Neurosciences Institute Interdisciplinary Scholar Award (C.N.B.), T32 AG000266 (C.N.B.), K99AG07849901 (C.N.B.), T32 AG0047126 (R.D.N.), the Iqbal Farrukh & Asad Jamal Center for Cognitive Health in Aging (R.D.N.), a Brain Resilience Scholar Award from the Knight Initiative for Brain Resilience at the Wu Tsai Neurosciences Institute (R.D.N.), and K99AG07687901 (R.D.N.).

## Author contributions

C.N.B., R.D.N., A.B., and K.D. designed the project; C.N.B. and R.D.N. built the tracking setup; C.N.B. and R.D.N. performed all experiments; C.N.B. performed all analysis with input from all authors; L.Z. and S.W.L. helped conceptualize tensor factorization and aging trajectory analyses, and developed software for syllable identification and hidden Markov modeling; C.N.B., R.D.N., A.B., and K.D. wrote the manuscript with input from L.Z. and S.W.L.; and K.D. and A.B. supervised all aspects of the project.

## Data and materials availability

Processed HMM data (10 min binning) and processed data resulting from TCA analysis are deposited at (https://doi.org/10.5281/zenodo.17238217) and available without restrictions upon publication. Processed counts for RNA-seq data generated in this study are deposited at (https://doi.org/10.5281/zenodo.17238217) and raw data will be deposited at NCBI-GEO for publication. Code for life-long behavioral analysis and RNA sequencing analysis are available at (https://doi.org/10.5281/zenodo.17371640). Code for custom JAX implementation of stochastic EM for hidden Markov models is available at https://github.com/lindermanlab/scalable-gaussian-hmm.

## Supplementary Materials

Figs. S1 to S19

Movies S1 to S2

Data S1 to S2

## Supplementary Materials

**Fig. S1.**
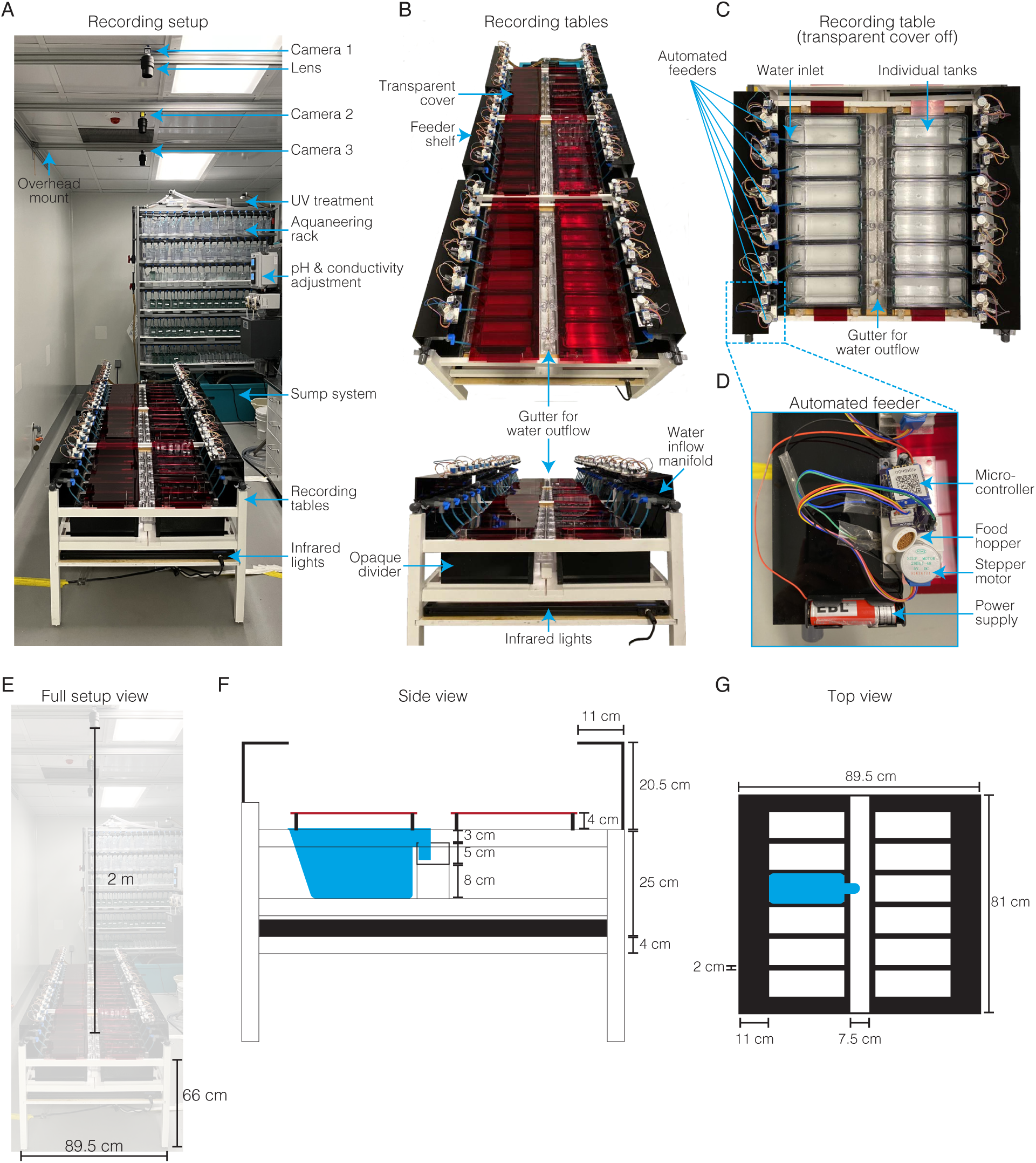
Long-term recording setup. (**A**) Three custom table racks with continuous water inflow and outflow from Aquaneering rack and sump system. Three overhead mounted infrared-sensitive cameras (one per table). (**B**) Twelve 2.8-L tanks fixed in each custom table rack. Infrared-transparent red acrylic lid covers tanks. Automated feeders mounted on a black acrylic shelf drops food into tanks at fixed times each day. Gutter collects tank outflow for filtering and recirculation through sump system. Water inflow manifold fixed to table racks with inlet valves and inlet tubes for each tank. Opaque black acrylic dividers between tanks prevents fish in adjacent tanks from seeing one another. Infrared backlight below tanks. (**C**) Overhead view of single table rack without acrylic cover. (**D**) Zoom in on automated feeder highlighted in (**C**) showing feeder components. (**E**-**G**) Measurements of the setup.

**Fig. S2.**
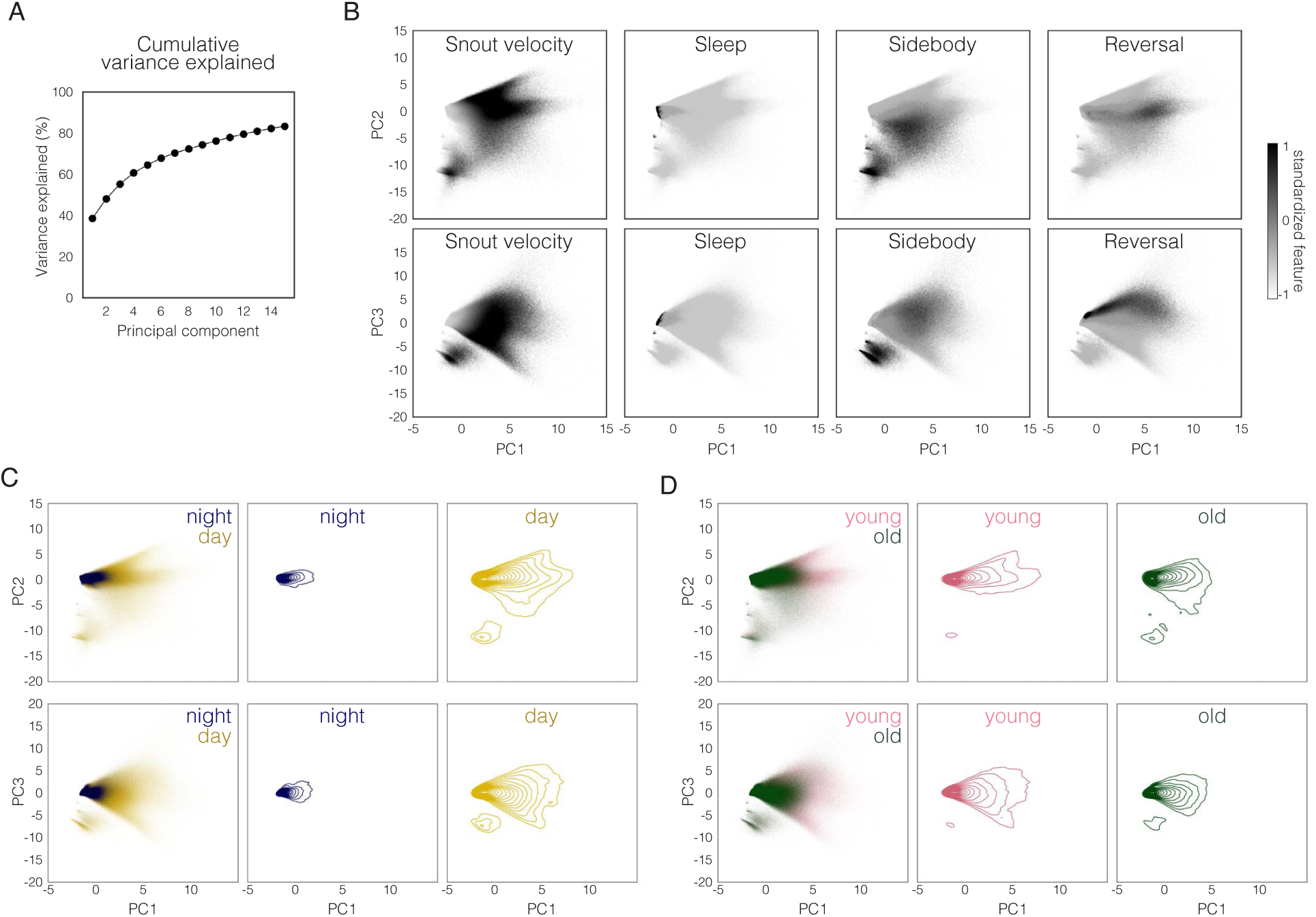
PCA of pose features. (**A**) Cumulative variance explained by up to 15 principal components used for training HMMs. (**B**) Top three pose feature PCs (PC1, PC2, PC3) of data from a single day (down-sampled from 20 Hz to 4 Hz frame rate) for nine different fish at two ages (45 days old and 270 days old). Each scatter point represents a single frame of recording and color maps indicating heatmap for four select pose features. (**C**) Same as (**B**) but colored by day (yellow) vs. night (blue) frames (left) and contour plots of frames separating day and night (right). (**D**) Same as (**B**) but colored by colored by young (pink: 45 days old) vs. old (green: 270 days old) frames (left) and contour plots of frames separating young and old (right).

**Fig. S3.**
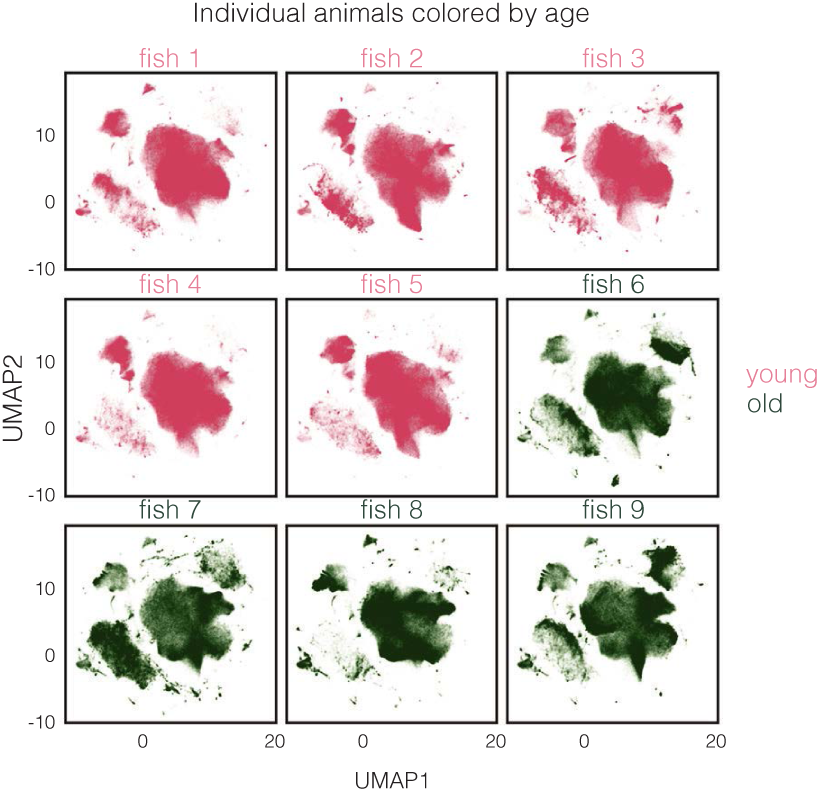
Individual animal pose features. UMAP embedding of pose features PCs of data from a single day (down-sampled from 20 Hz to 4 Hz frame rate) for nine different fish at two ages (pink: 45 days old, young; and green: 270 days old, old). Each scatter point represents the embedding of a single frame of recording.

**Fig. S4.**
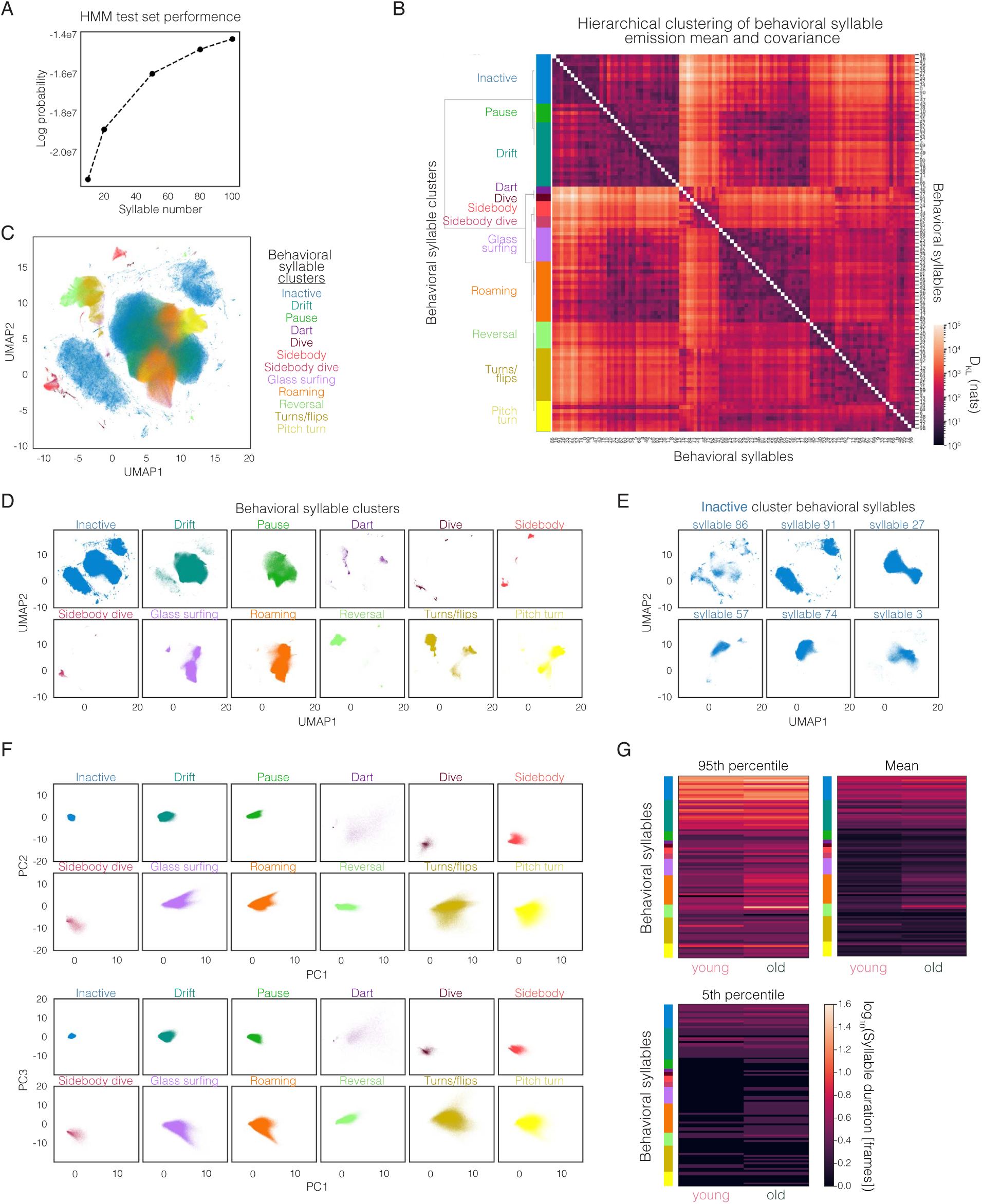
HMM optimization and HMM-derived behavioral syllable clustering. (**A**) Cross-validation log probability of different HMM models with varying numbers of behavioral syllables using stochastic EM fitting procedure. (**B**) Hierarchical clustering of behavioral syllables using symmetrized D_KL_ of HMM emission distributions with the resulting behavioral syllable clusters shown in color along left. (**C**) UMAP embedding of pose features PCs of data from a single day for eight different fish (five young and four old) down-sampled from 20 Hz to 4 Hz frame rate, with each scatter point representing the embedding of a single frame of recording and color maps indicating the behavioral syllable cluster. (**D**) Same as (**C**) but frames from distinct behavioral syllable clusters are plotted separately. (**E**) Same as (**C**) but only plotting frames from distinct behavioral syllables within the “inactive” behavioral syllable cluster. (**F**) Top three pose feature PCs (PC1, PC2, PC3) with color maps indicating the behavioral syllable cluster and distinct behavioral syllable clusters plotted separately. (**G**) Mean, 5^th^ percentile, and 95^th^ percentile syllable duration (frames) comparing young (45-day old) and old (270-day old) animals from a single day for nine different fish (five young and four old).

**Fig. S5.**
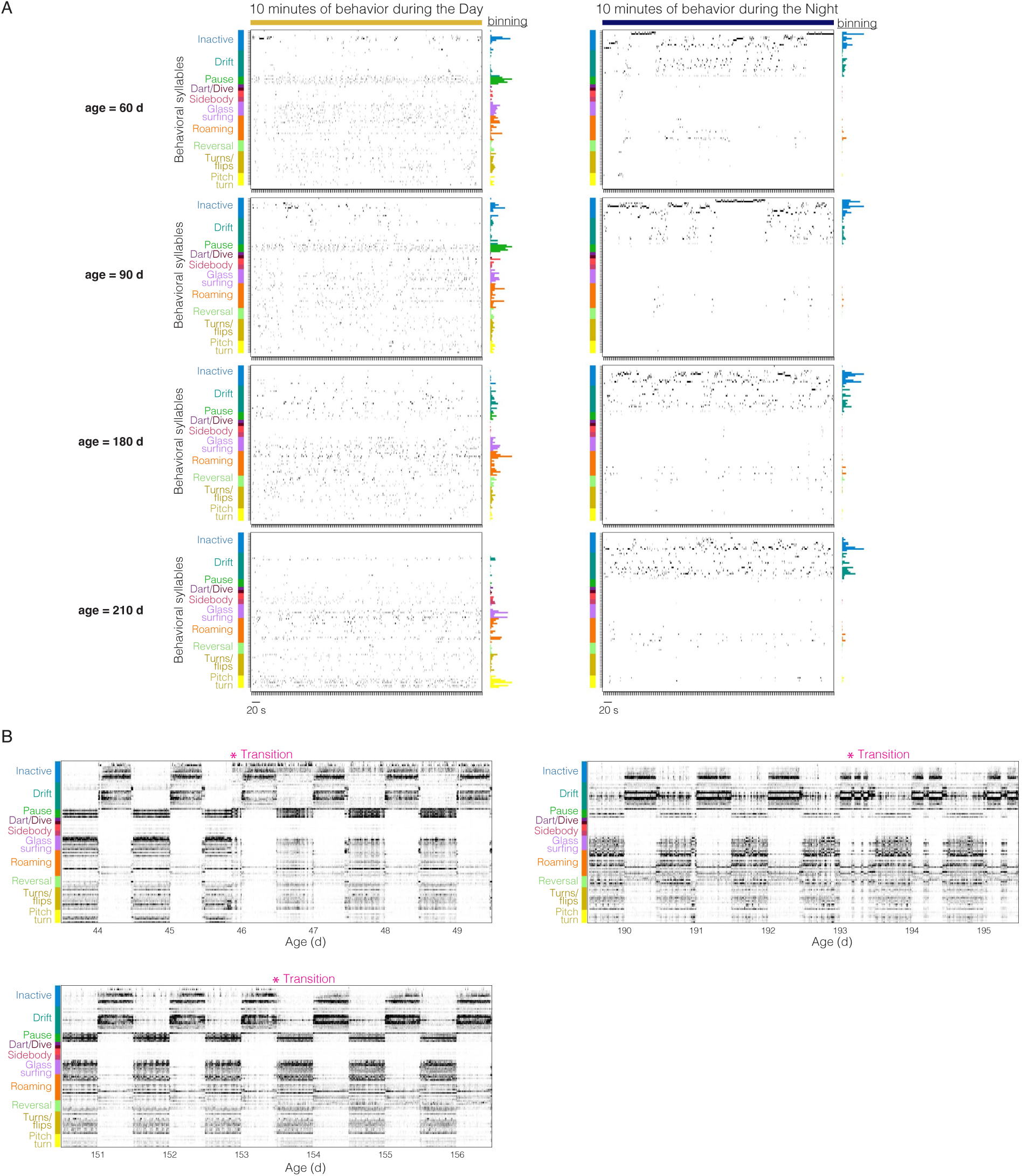
Behavioral syllable dynamics at different ages and during transitions. (**A**) Sampling of 100 HMM-derived behavioral syllables for a single fish at four different ages during 10-min interval during the day (left) and 10-min interval during the night (right). Throughout figure, behavioral syllables are ordered according to similarity and hierarchical clustering of distinct types of behavior shown in color along left. Behavioral syllables are ordered as in Fig. 1I. Histogram shows time spent in each behavioral syllable during the 10-min bin on right. (**B**) Behavioral syllable usage for a single animal for windows of life during a transition highlighted in Fig. 1K.

**Fig. S6.**
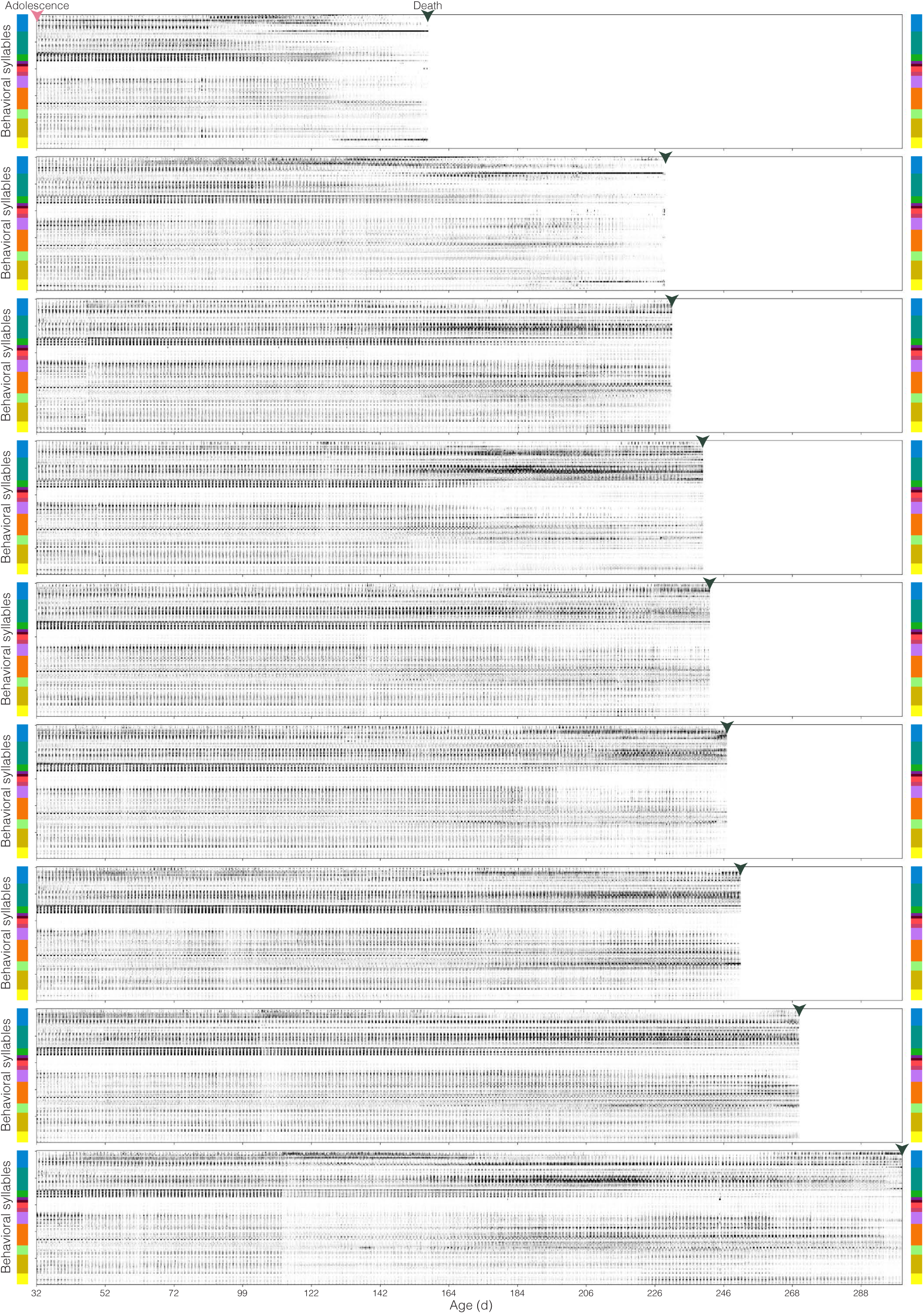
Whole-lifespan behavioral syllable use for individual animals from adolescence until death. Animals ordered based on lifespan from shortest to longest lived. Behavioral syllables are ordered according to similarity and hierarchical clustering of distinct types of behavior shown in color along left and right. Behavioral syllables are ordered as in Fig. 1I.

**Fig. S7.**
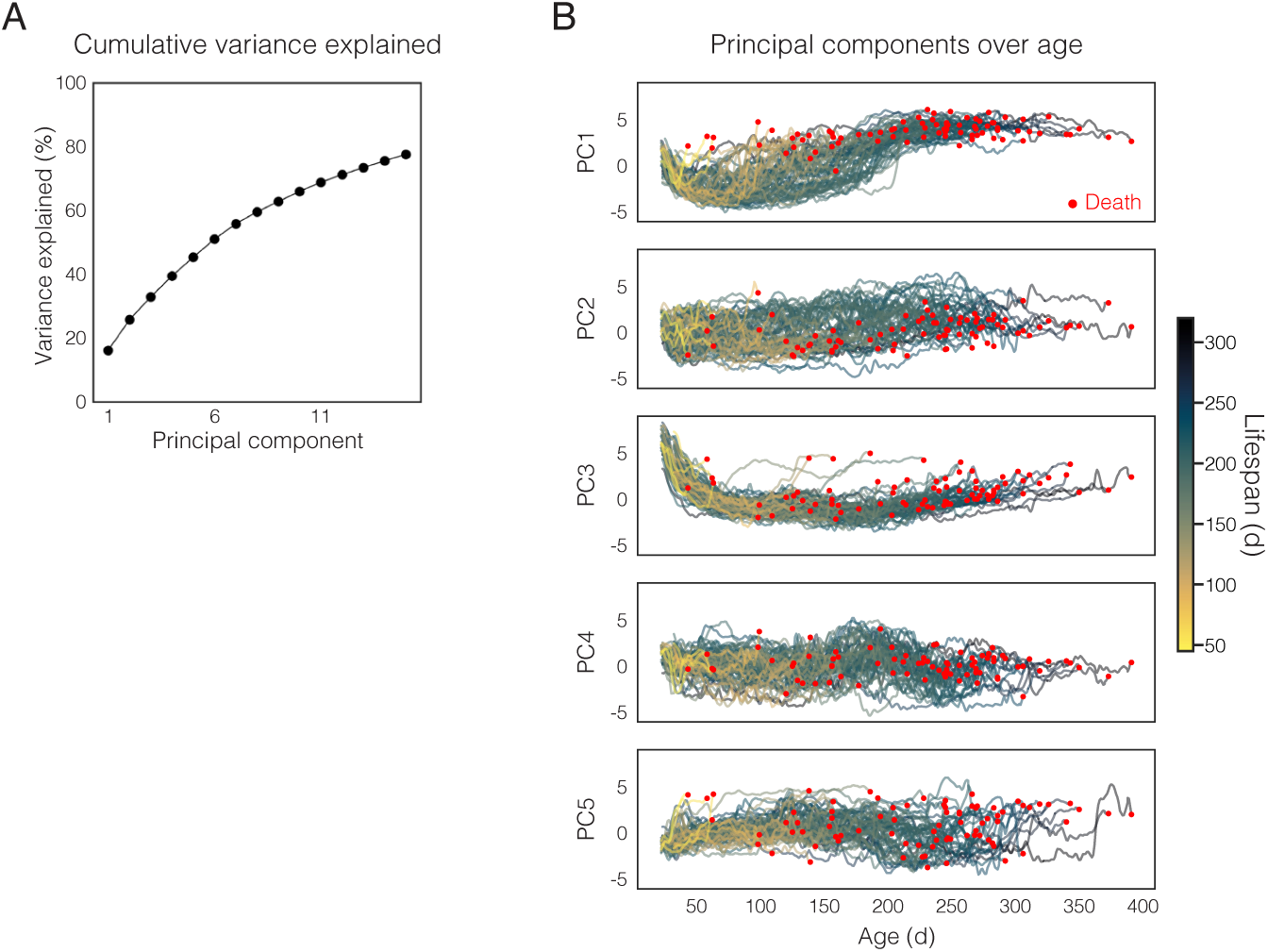
PCA of 45 tensor components. (**A**) Cumulative variance explained for increasing principal components. The top three PCs are used in visualizations in Fig. 2E-G and Fig. 3B,H. (**B**) Top five PCs plotted across age of 81 animals colored by future lifespan. Age of death indicated by red circle for each animal.

**Fig. S8.**
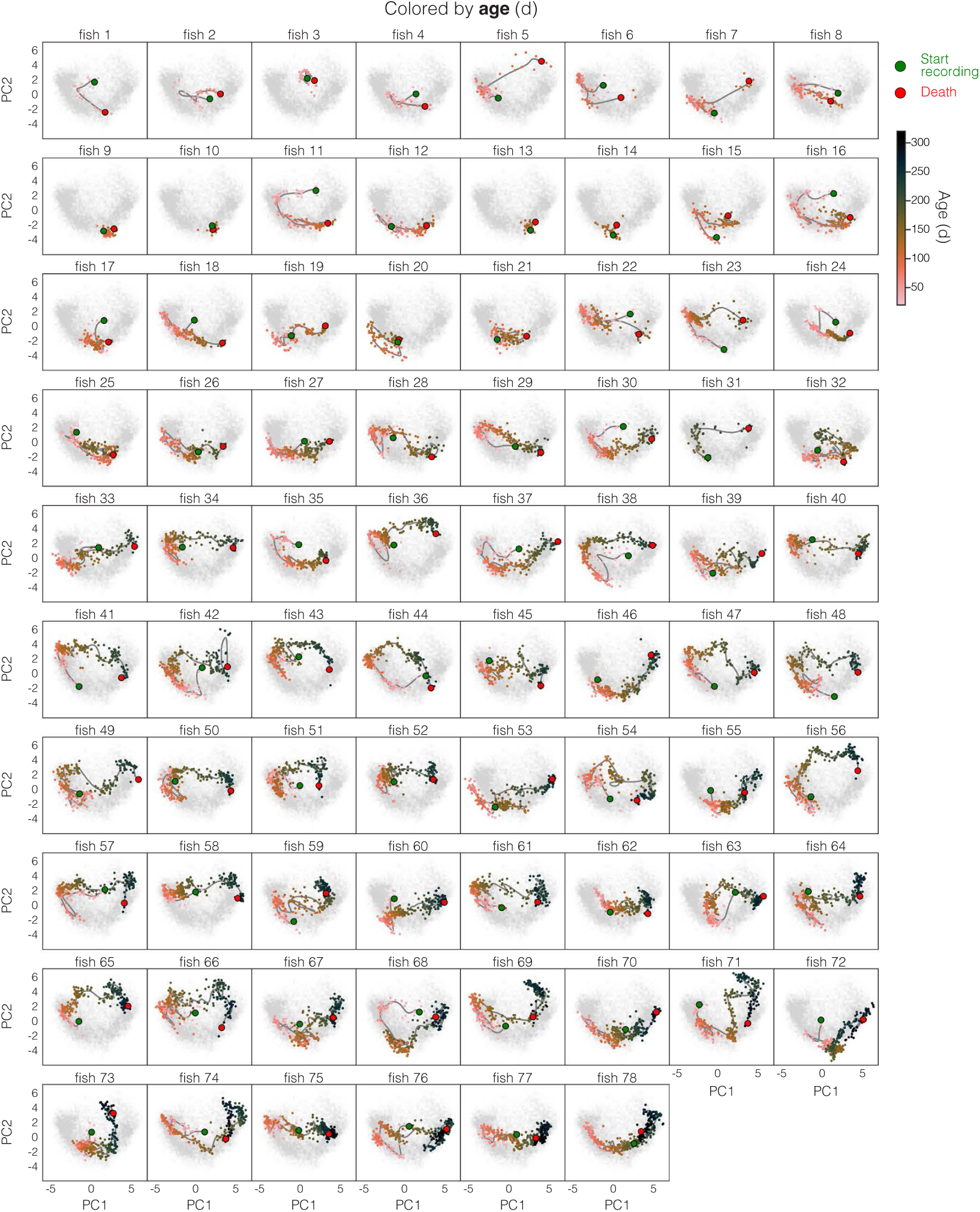
Individual animal aging trajectories. Top two PCs (PC1 and PC2) of 45 TC data to visualize aging trajectories of 78 individual animals from start of recording (green circle) to death (red circle). Each animal is shown in a separate plot. Points show individual days. Line shows smoothed trajectory. Each point colored by animal age. The whole population is shown in light grey scatter points for reference.

**Fig. S9.**
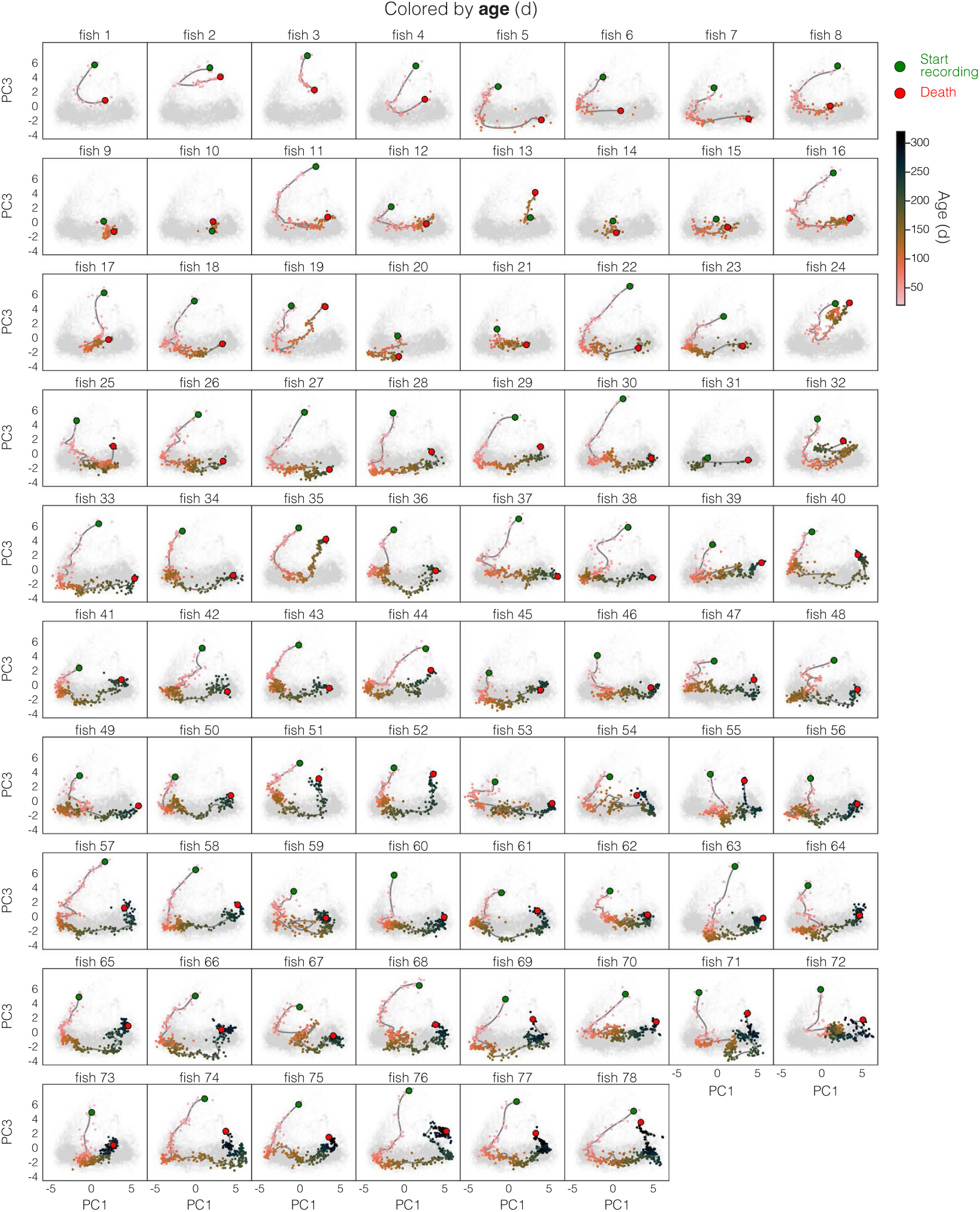
Individual animal aging trajectories. Top three PCs (PC1 and PC3) of 45 TC data to visualize aging trajectories of 78 individual animals from start of recording (green circle) to death (red circle). Each animal is shown in a separate plot. Points show individual days. Line shows smoothed trajectory. Each point colored by animal age. The whole population is shown in light grey scatter points for reference.

**Fig. S10.**
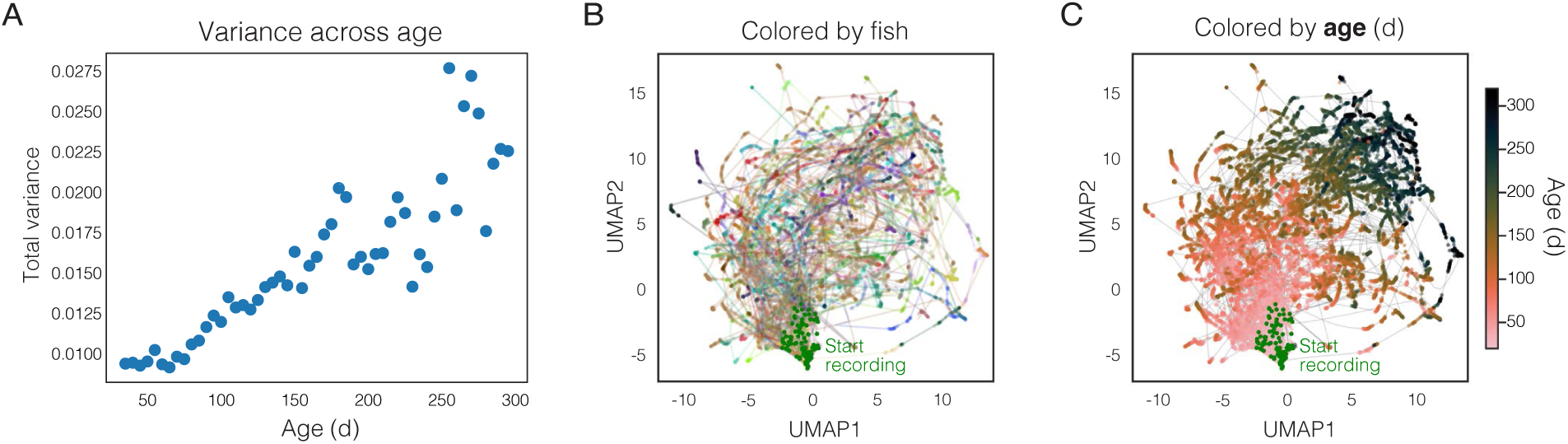
Aging trajectories across life. (**A**) Summed variance of all 45 TCs at a range of ages across life. (**B**-**C**) UMAP embedding of 45 TC data to visualize aging trajectories. Points show individual days. Line shows smoothed trajectory. With each animal in a different color (**B**) or with each point colored by animal age (**C**).

**Fig. S11.**
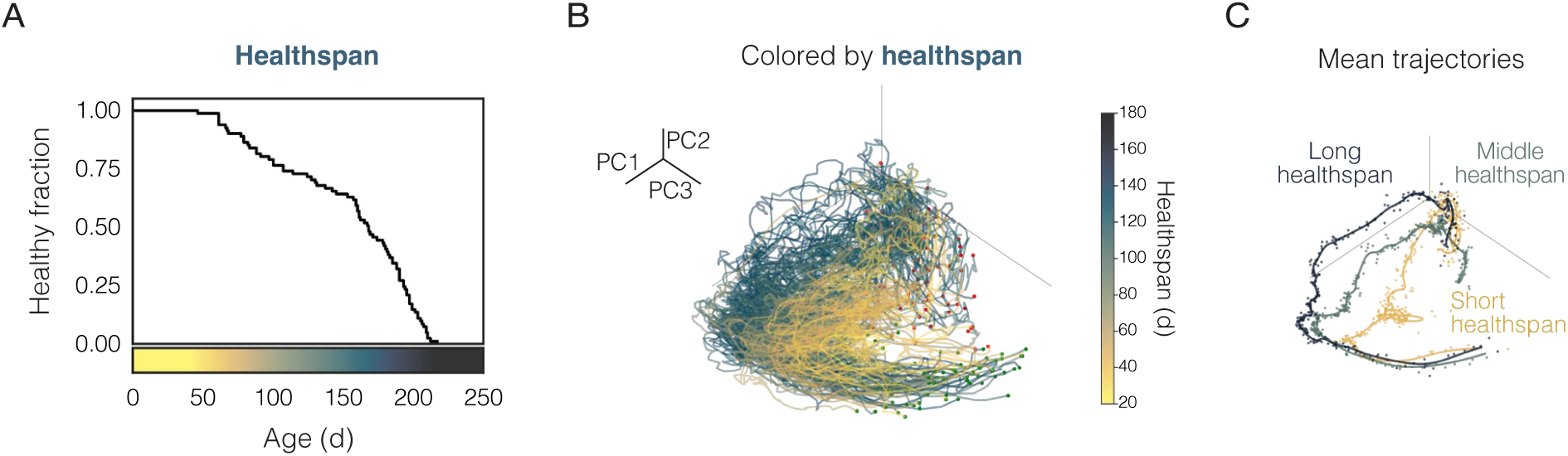
**Behavior-based aging trajectories of short-healthspan vs. long-healthspan animals differ**. (**A**) Kaplan–Meier healthspan curve of tracked animals (n=81). (**B**) PCA aging trajectories of 81 animals colored by healthspan (i.e., healthy life), defined for killifish as remaining active and consuming all available food. (**C**) Mean trajectory of distinct healthspan groups (short: 25^th^ percentile; middle: 25-50^th^ percentile; long: 75^th^ percentile).

**Fig. S12.**
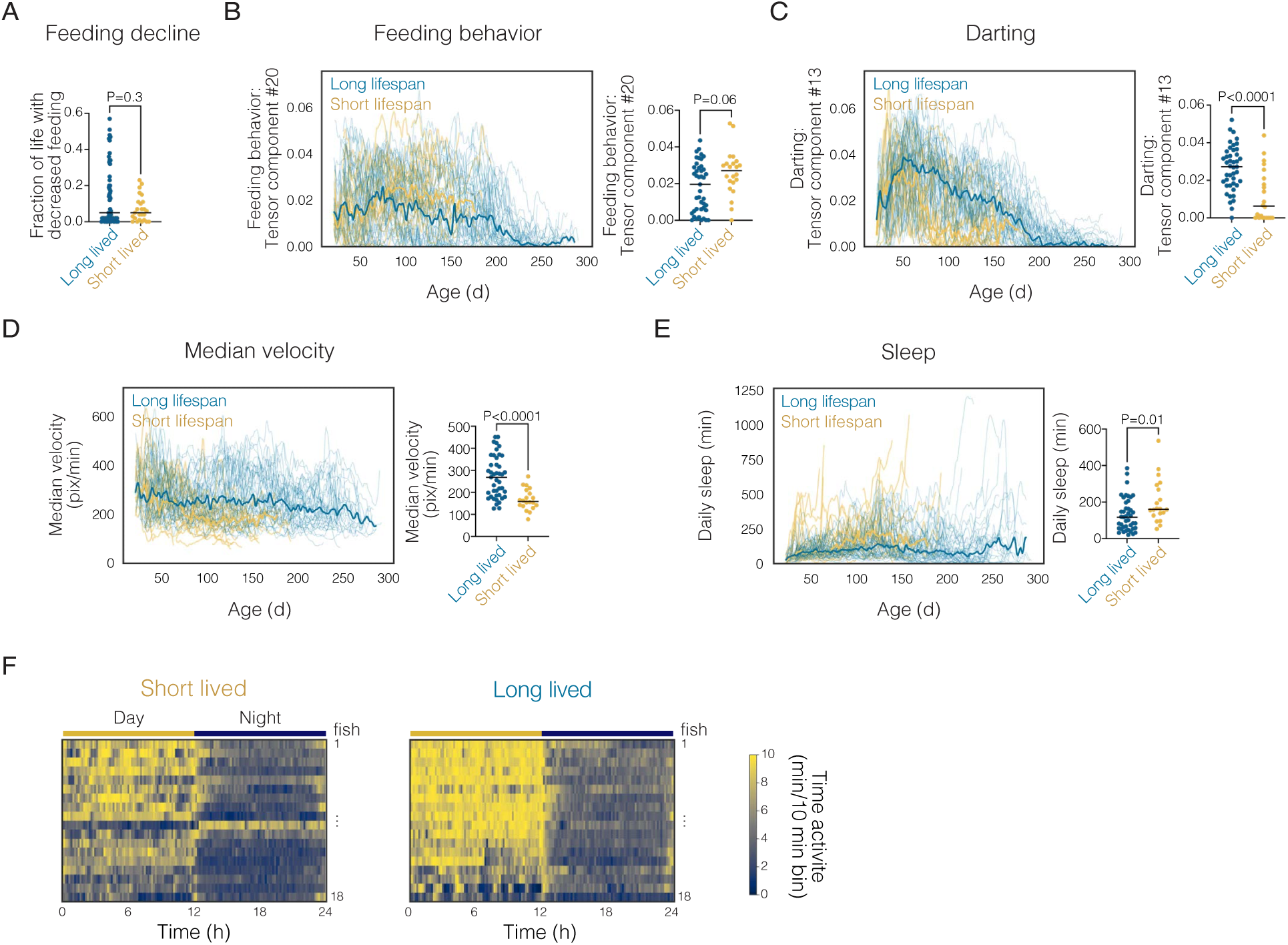
**Behavior and locomotor activity of long-vs. short-lived animals**. (**A**) Proportion of life prior to death with reduced food consumption resulting in residual uneaten food. Feeding behavior (weighting of tensor component #20 in Fig. 2D) (**B**), vigorous darting behavior (weighting of tensor component #13 in Fig. 2D) (**C**), median velocity (**D**), and total daily sleep (**E**) across life (left) and at 100 days old (right) grouped by lifespan. Mann Whitney test for significances. (**F**) Heatmap of time spent active across 24 hr at 100 days old with each row representing a different animal and animals grouped by lifespan. Light/dark cycle indicated by yellow/blue bar.

**Fig. S13.**
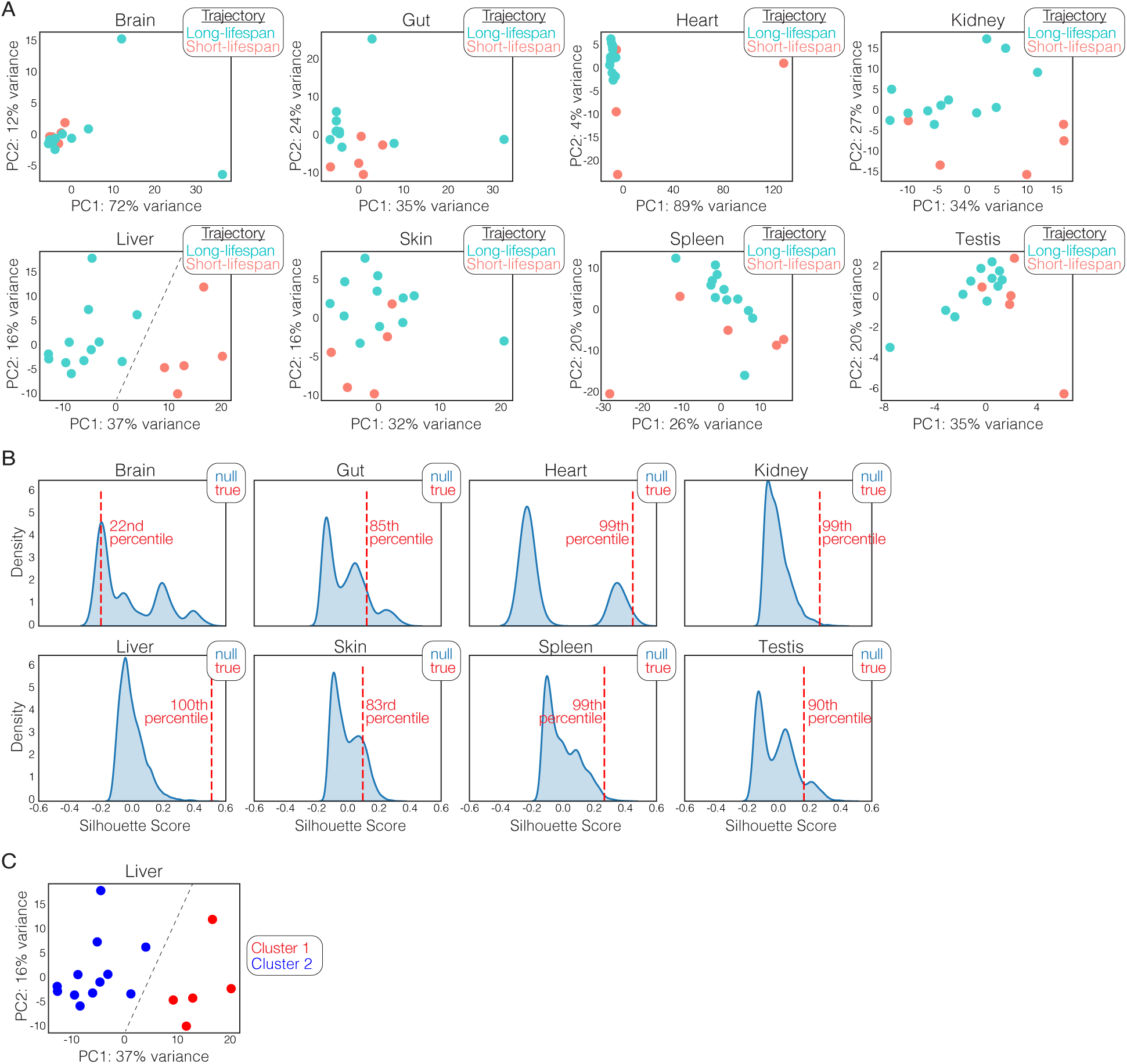
**Multi-organ transcriptomics**. (**A**) PCA of whole transcriptome for each animal in the middle-aged group (150 d; n=17) colored by future-lifespan trajectory based on behavior. Each plot is a different organ. (**B**) Silhouette score in PC1/PC2 space of true labels (long-lifespan vs short-lifespan trajectory based on behavior) indicated by red dashed line and calculated percentile compared to null distribution build by permutations of randomized group assignment across all n=17 samples (blue distribution). (**C**) Unbiased k-means clustering of the liver PC1/PC2 data into two clusters results in perfect match of the observed short-lifespan and long-lifespan trajectory based on behavior labels shown in (**A**).

**Fig. S14.**
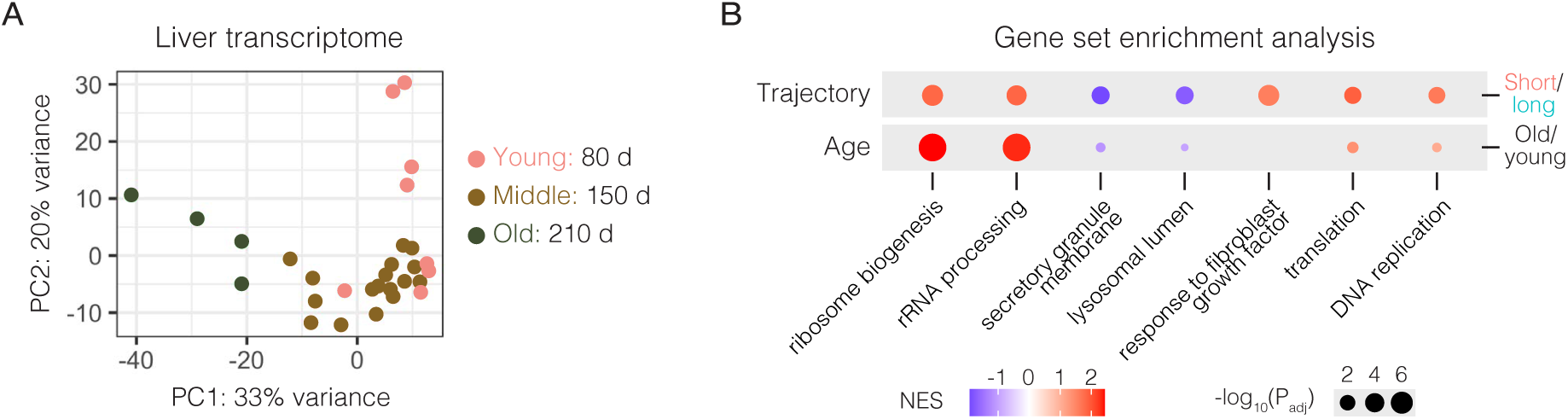
**Liver transcriptome is different both with age and lifespan trajectory**. (**A**) PCA of whole transcriptome for each animal combining the young (80 d; n=8), middle-aged (150 d; n=17), and old (≥210 d, n=4) cohorts colored age. (**B**) Gene set enrichment analysis of the liver transcriptome as a function of lifespan trajectories based on behavior (short-vs. long-lifespan trajectory) compared to age (old vs. young) highlighting significant Gene Ontology (GO) terms. NES, normalized enrichment score. Padj, P-value adjusted for multiple hypothesis testing

**Fig. S15.**
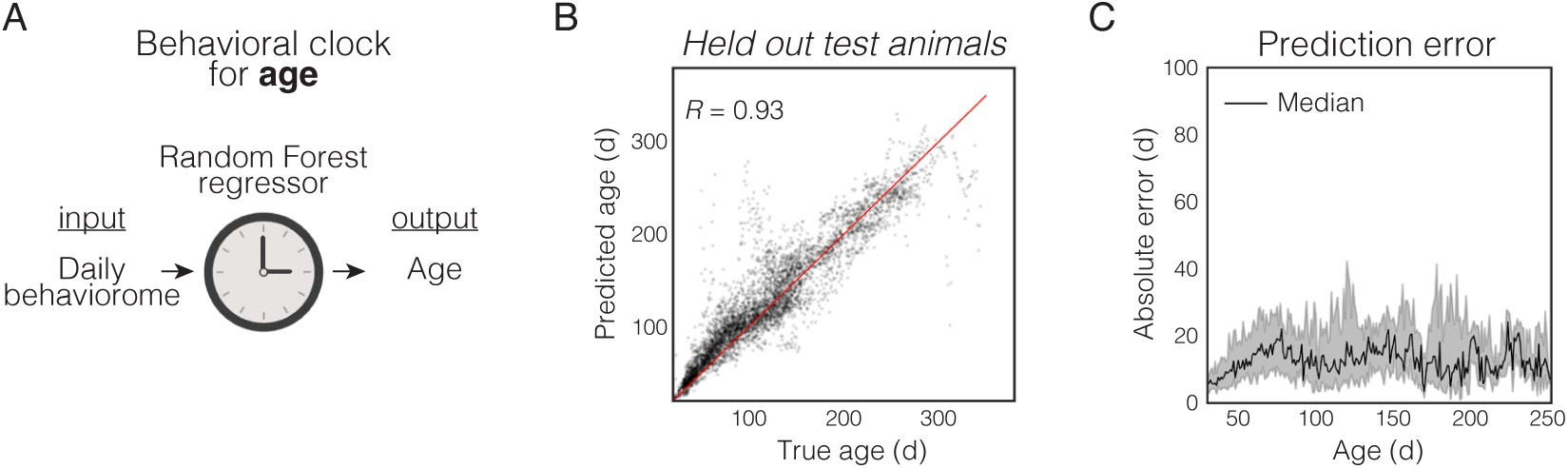
Behavioral clock for age. (**A**) Random forest regression models trained on behavioral TC data to predict age (i.e., behavioral clock for age). (**B**) Held out test set cohorts (36 held-out animals) model prediction vs. true age. (**C**) Held out test set cohorts (36 held-out animals) model prediction absolute error calculated across age.

**Fig. S16.**
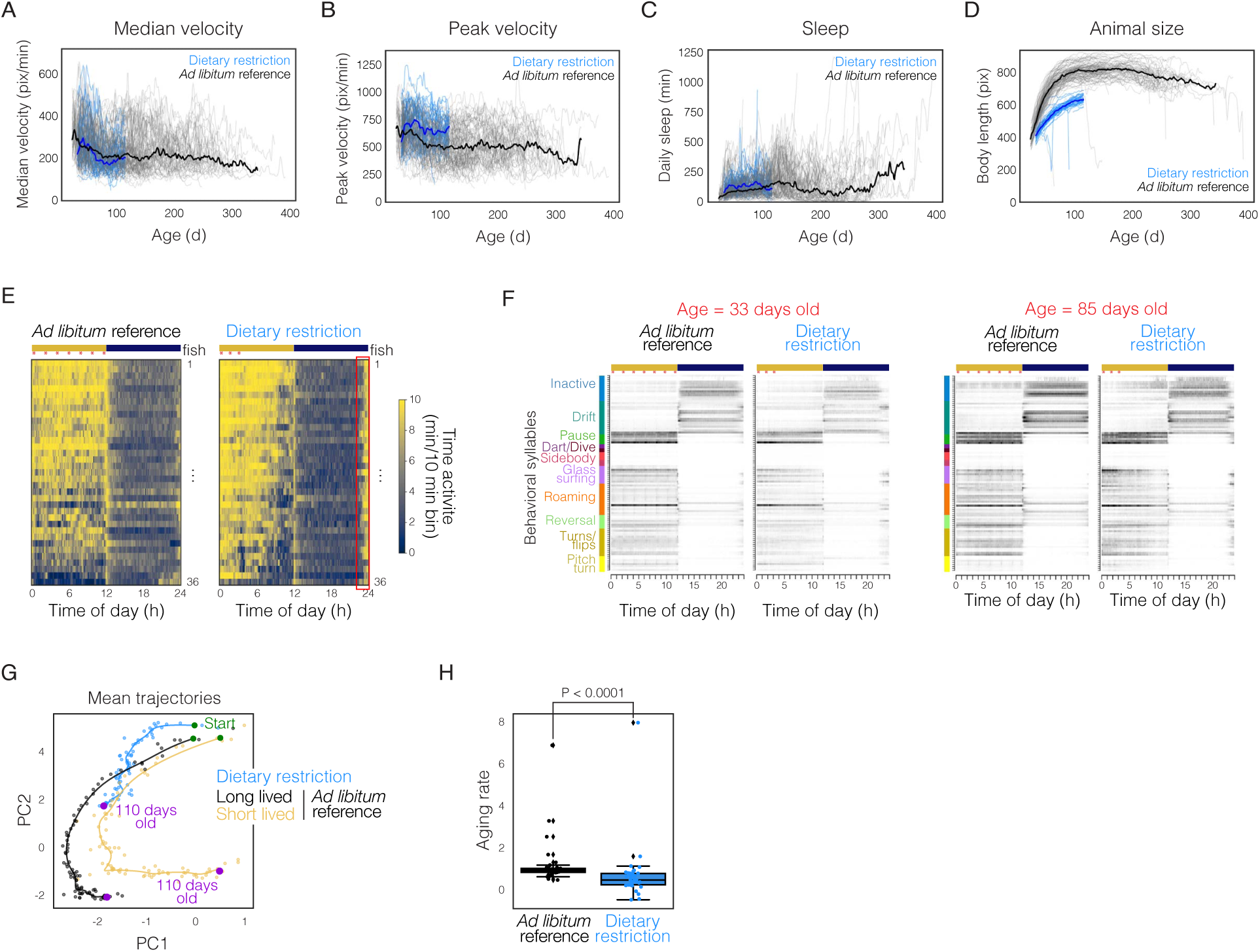
**Behavior, locomotor activity, and size with dietary restricted feeding**. Median velocity (**A**), peak velocity (**B**), total daily sleep (**C**), and animal size (**D**) across recordings comparing animals under dietary restriction (n=39; recorded until 110 d old) to *ad libitum* reference (n=80; recorded full lifespan). (**E**) Heatmap of time spent active across 24 hr at 54 d old with each row representing a different animal. Animals under dietary restriction (right; n=36) are compared to randomly selected *ad libitum* reference animals from previous cohorts (left; n=36). Red asterisks indicate feeding times. Light/dark cycle is indicated by yellow/blue bar. Red rectangle underscores elevated activity before the lights turn on for the animals under dietary restriction. (**F**) Mean behavioral syllable usage across 24 hr for all animals under dietary restriction (right; n=39) and all animals from the *ad libitum* reference cohorts (left; n=80) at 33 d old and 85 d old. Behavioral syllables are ordered as in Fig. 1I. (**G**) Mean behavioral aging trajectory of long-lived *ad libitum* reference animals (black; lifespan > 200 days); short-lived *ad libitum* reference animals (yellow; lifespan<200 days); vs. dietary restriction animals (blue). Purple points indicate when animals reach 110 days old. By 110 days, the dietary restriction animals are behaviorally more similar to ∼50-day old *ad libitum* animals. However, it is not possible to determine if the dietary restricted animals follow an *ad libitum* short-lived or long-lived trajectory because at this point in the trajectory, there is not yet a robust difference between the behavior of short-lived and long-lived animals (Fig. 3B,D-E; Fig. 4F-G,I). (**H**) Aging rate (i.e., the slope of behavioral clock predictions vs true age) of *ad libitum* reference animals (black) vs. dietary restriction animals (blue).

**Fig. S17.**
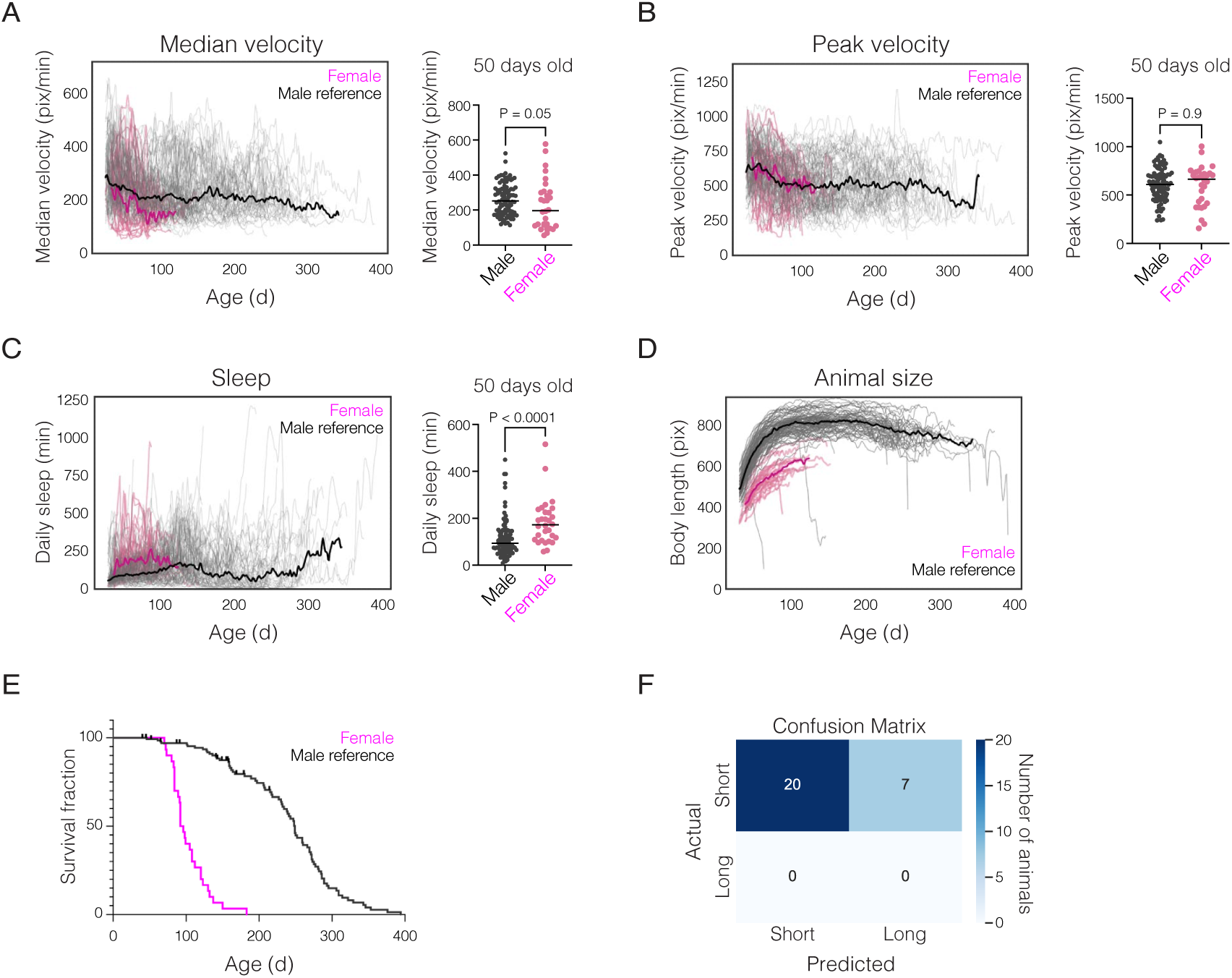
**Behavior, locomotor activity, size and lifespan of females**. Median velocity (**A**), peak velocity (**B**), and total daily sleep (**C**) across life (left) and at 50 days old (right) comparing females to the male reference. Mann Whitney test for significance. (**D**) Animal size across life. (E) Kaplan–Meier lifespan curve of female animals (n=31) compared to male reference (n=118). (F) Confusion matrix for random forest classification model predictions on the female cohort (n=27) based on behavior at 70 days old. Animals are classified as either short-or long-lived (i.e., behavioral classifier for lifespan).

**Fig. S18.**
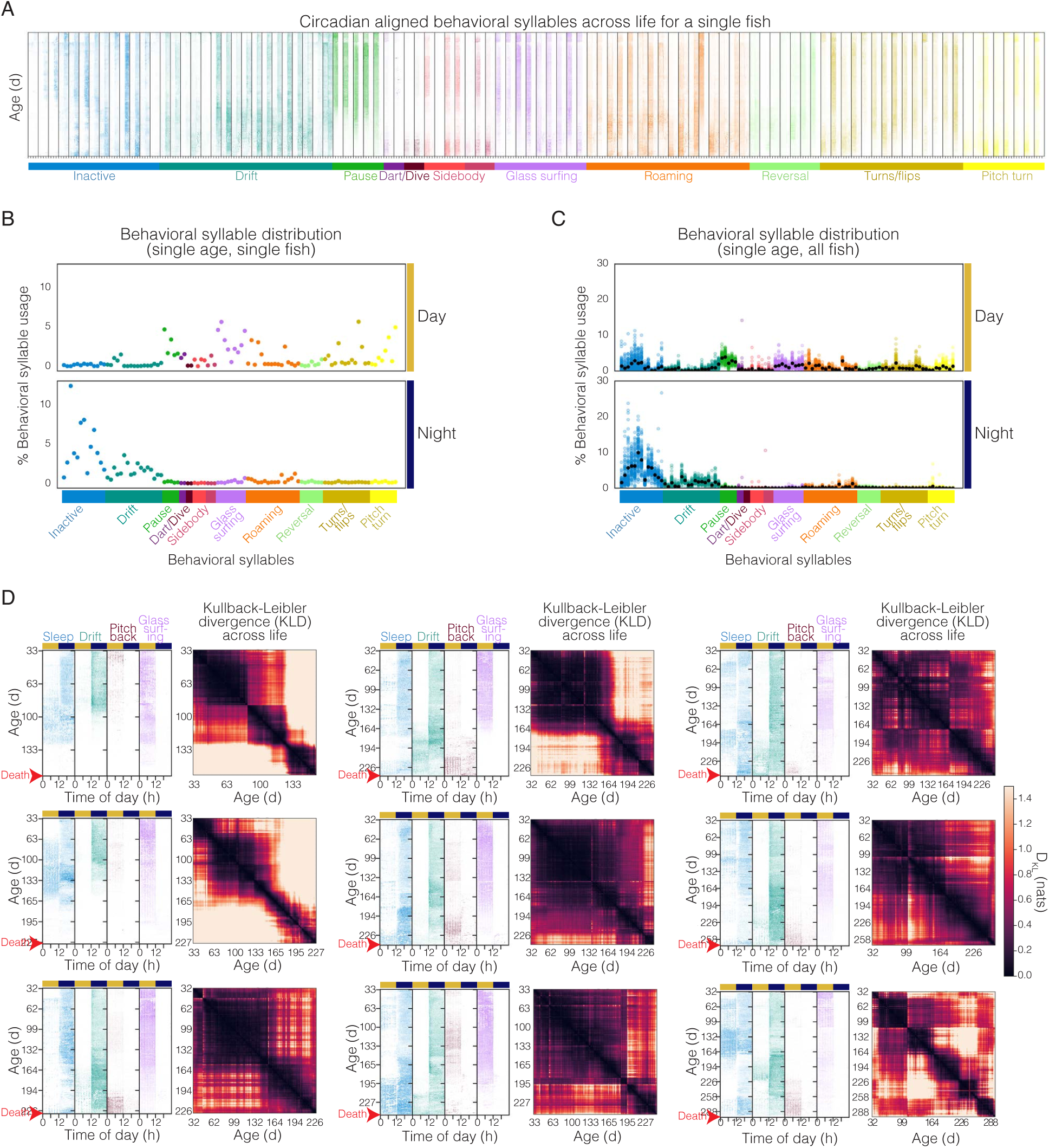
Changes in whole-lifespan behavioral syllable use for individual animals. (**A**) Circadian-aligned, whole lifespan heatmaps for all 100 behavioral syllables of example animal in Fig. 6A-D. (**B**) Behavioral syllable usage distribution for day and night of a single animal at a single age. (**C**) Behavioral syllable usage distribution for day and night of all tracked animal at a single age. (**D**) Circadian-aligned, whole-lifespan heatmaps of four select behaviors (as in Fig. 6B) for nine individual animals (same animals shown in fig. S6) aligned with cross-correlation of symmetrized DKL of the behavioral syllable usage distribution comparing all days in one animal’s life to all other days ordered from youth (32 d) to death. Light/dark cycle is indicated by yellow/blue bar.

**Fig. S19.**
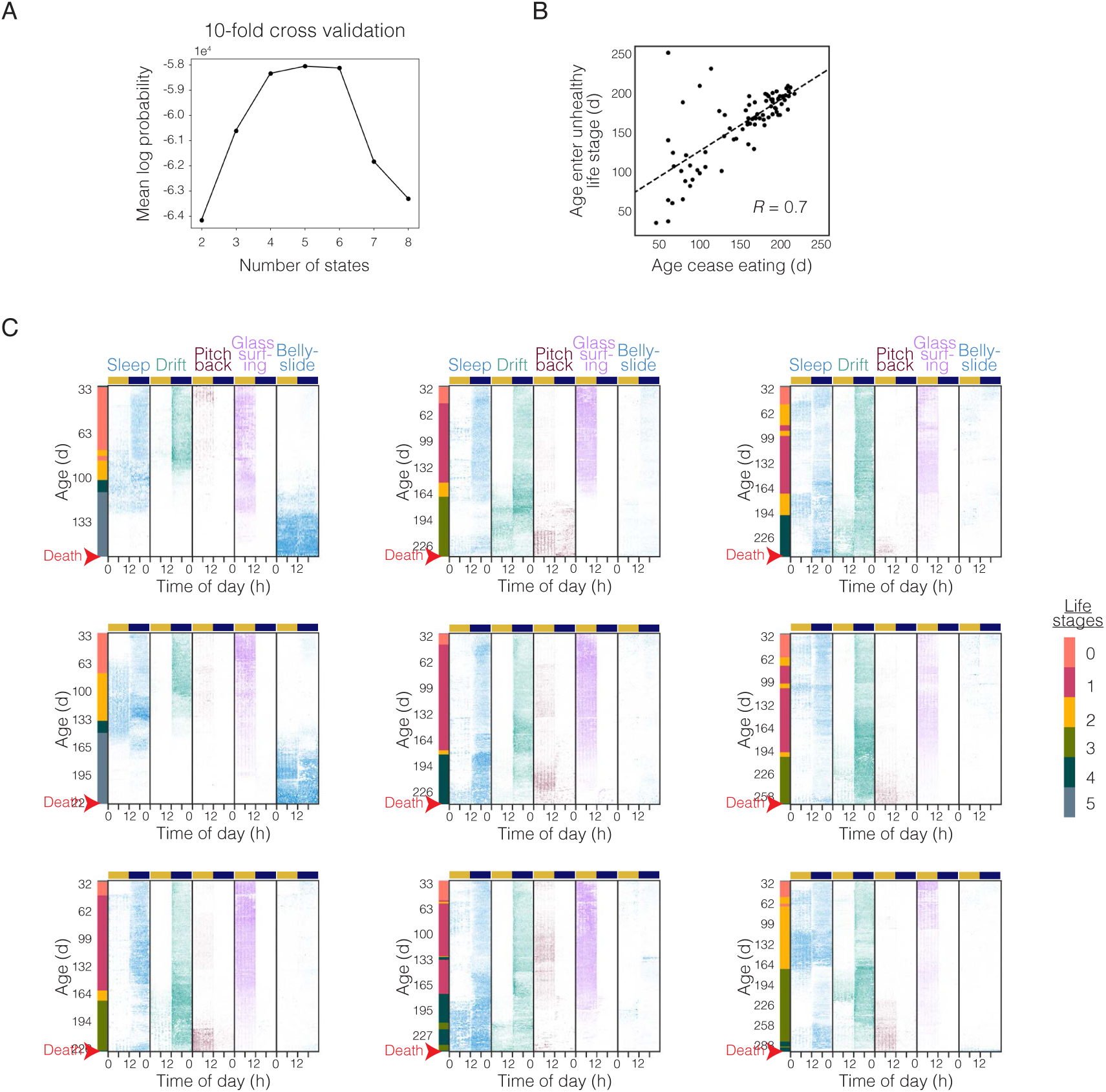
HMM life stage model optimization and output. (**A**) Mean log probability of 10-fold cross validation varying the number of states. (**B**) Correlation of age at which animals cease eating, with the youngest age an animal enters either unhealthy (green or blue) life stage. (**C**) Circadian-aligned, whole-lifespan heatmaps of five select behaviors for nine individual animals (same animals shown in fig. S6 and fig. S18) aligned with HMM-predicted life stages along the left. Light/dark cycle is indicated by yellow/blue bar.

**Movie S1.**

Example movie from one recording table of 12 male killifish with cameras mounted overhead with infrared backlighting (20 frames per second).

**Movie S2.**

Deep convolutional neural network prediction of key points along the body – snout, midbody, endbody, tail, fan, and sidebody for example Movie S1.

**Data S1. (separate file)**

Fifty-seven features describing pose feature dynamics across time calculated by either the mean and/or standard deviation (depending on the feature) of pose features (evaluated from the x-/y-coordinates of tracked key points) over a 10-frame rolling window.

**Data S2. (separate file)**

Liver transcriptomics comparison of short-lifespan vs long-lifespan trajectory groups at 150 days old (n=17) shown in Fig. 3G-K. (**DEseq sheet**) Differential expression analysis results with log2FoldChange>0 if gene expression is elevated in the short-lifespan trajectory group relative to the long-lifespan trajectory group. (**PEA sheet**) Over-representation analysis for significantly differentially expressed genes between short-lived vs. long-lived trajectory using Gene Ontology (GO) enrichment analysis. GO pathways are up (positive charge) if driven by genes significantly elevated in the short-lifespan trajectory group relative to the long-lifespan trajectory group. (**GSEA sheet**) Gene set enrichment analysis results with positive enrichment score if elevated in the short-lifespan trajectory group relative to the long-lifespan trajectory group. (**TopGenes sheet**) Differentially expressed genes (Padj > 0.1) between the short-lifespan vs long-lifespan trajectory groups at 150 days old grouped by pathways. Focusing on pathways elevated in the short-lifespan trajectory group relative to the long-lifespan trajectory group.

